# Undying leafcutter-ant colonies: The oldest *Atta texana* leafcutter-ant colonies are estimated to grow older than 100 years

**DOI:** 10.64898/2026.08.05.743090

**Authors:** Ulrich G. Mueller, Caroline E. Farrior

## Abstract

Tropical *Atta* leafcutter-ant colonies reach a maximum lifespan of 15-20 years because each colony has only a single queen, queens live for maximally 15-20 years, and colonies do not requeen after queen death. At the northern range limit of leafcutter ants in Louisiana, however, we found that the mortality rate of *Atta texana* colonies is consistent with a life expectancy of 65-75 years and a maximum colony lifespan exceeding 100 years. Annual mortality of established *A. texana* colonies in Louisiana is only 1-2% compared to 10-25% in tropical *Atta* species. Exceptional longevity of *A. texana* colonies is due to (i) absence of *Nomamyrmex* army-ant predators specialized to raid *Atta* colonies; (ii) polygyny of *A. texana* colonies, creating a unique social organization in *A. texana* colonies that differs from the monogynous colonies typical for all other *Atta* species; and (iii) adoption of young queens by old colonies, possibly facilitated by low genetic diversity in range-limit populations of *A. texana*. Centenarian longevity of *A. texana* colonies was so far unknown because, from the 1930s to the turn of the century, research prioritized eradication of *A. texana* as a forestry pest, and landscape-wide pesticide application by aircraft eliminated old *A. texana* colonies. After the most harmful pesticides were banned 30-50 years ago, *A. texana* populations recovered, yet *A. texana* required decades to regain its earlier niche in the forest ecology. In undisturbed habitat, well-established old *A. texana* colonies with large territories appear to recruit young queens especially well. Old age of a colony therefore does not necessarily increase mortality as in typical lifespan-limiting processes where age drives senescence and mortality (old age → mortality); instead, well-rooted residency that is correlated to age of an *A. texana* colony increases the chance of rejuvenation via queen adoption, reduces or eliminates reproductive senescence of a colony, and consequently prolongs colony lifespan (well-established residency → rejuvenation → long life). Large size attained in old age imparts potentially unending life.

## Introduction

Leafcutter ant colonies in the genus *Atta* are among the longest-living ant colonies because of an extraordinary longevity of *Atta* queens (KELLER 1998, COLE 2009, DA SILVA 2022, JAIMES-NINO & OETTLER 2025). Under laboratory conditions, for example, *Atta* queens have been observed to live up to 15-20 years (Table 1). Following queen death, an *Atta* laboratory colony deteriorates within months (AUTUORI 1950a, MARICONI 1970, WEBER 1982, WEISS 1990), and re-queening attempts of queenless colonies of tropical *Atta* species have been largely unsuccessful (EIDMANN 1936, MARICONI 1970, WEBER 1972, MOSER 1984, FOWLER & al. 1986, U.G. Mueller, unpubl.), so the lifespan of *Atta* queens in laboratory colonies predicts a corresponding lifespan of field colonies of 15-20 years or less. This prediction has been confirmed in multiple field surveys, which documented a maximum lifespan of 10-20 years for *Atta* colonies in the tropical range of leafcutter ants (Uruguay/Argentina to USA) (Table 1).

**Table 1.**
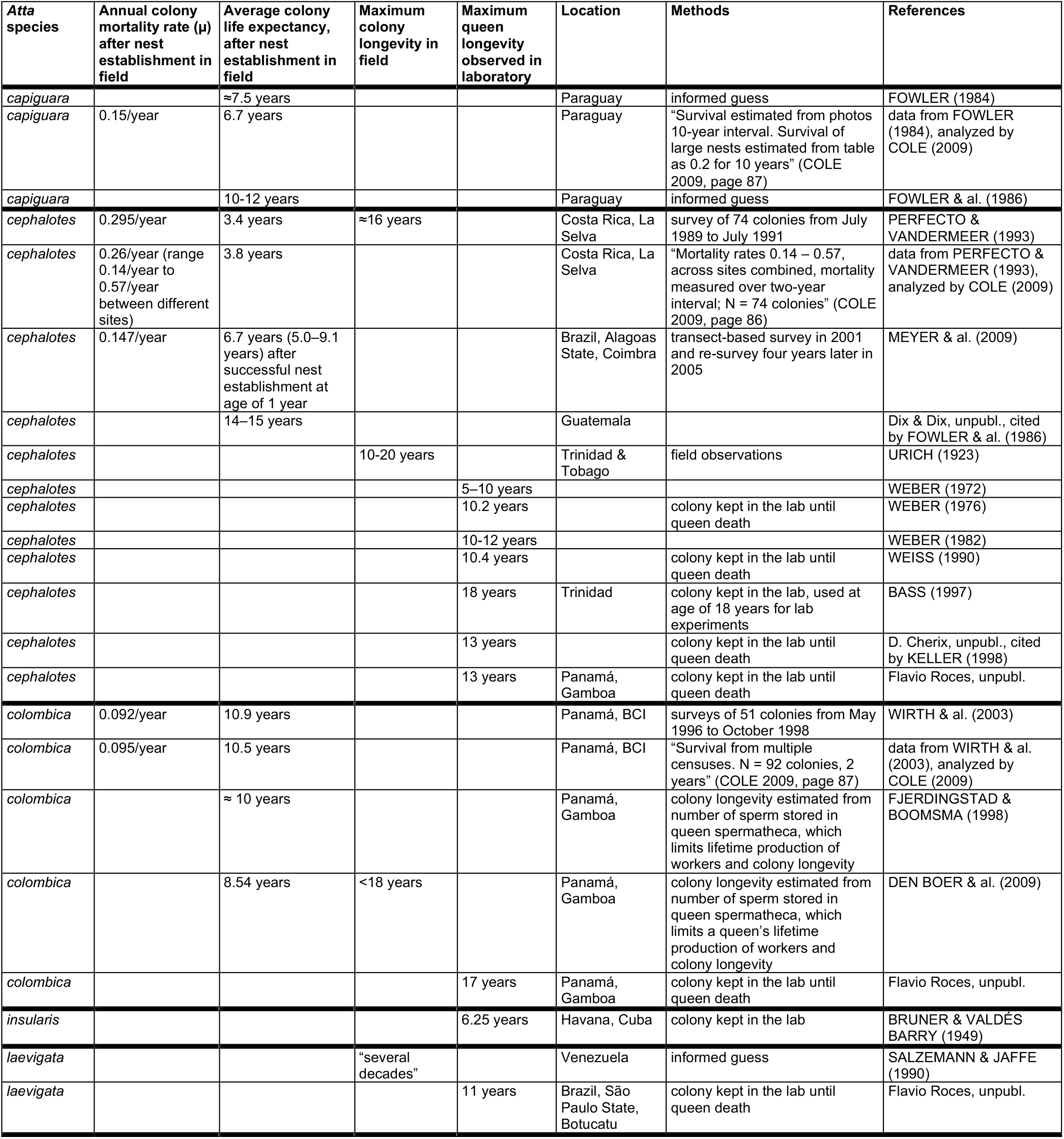
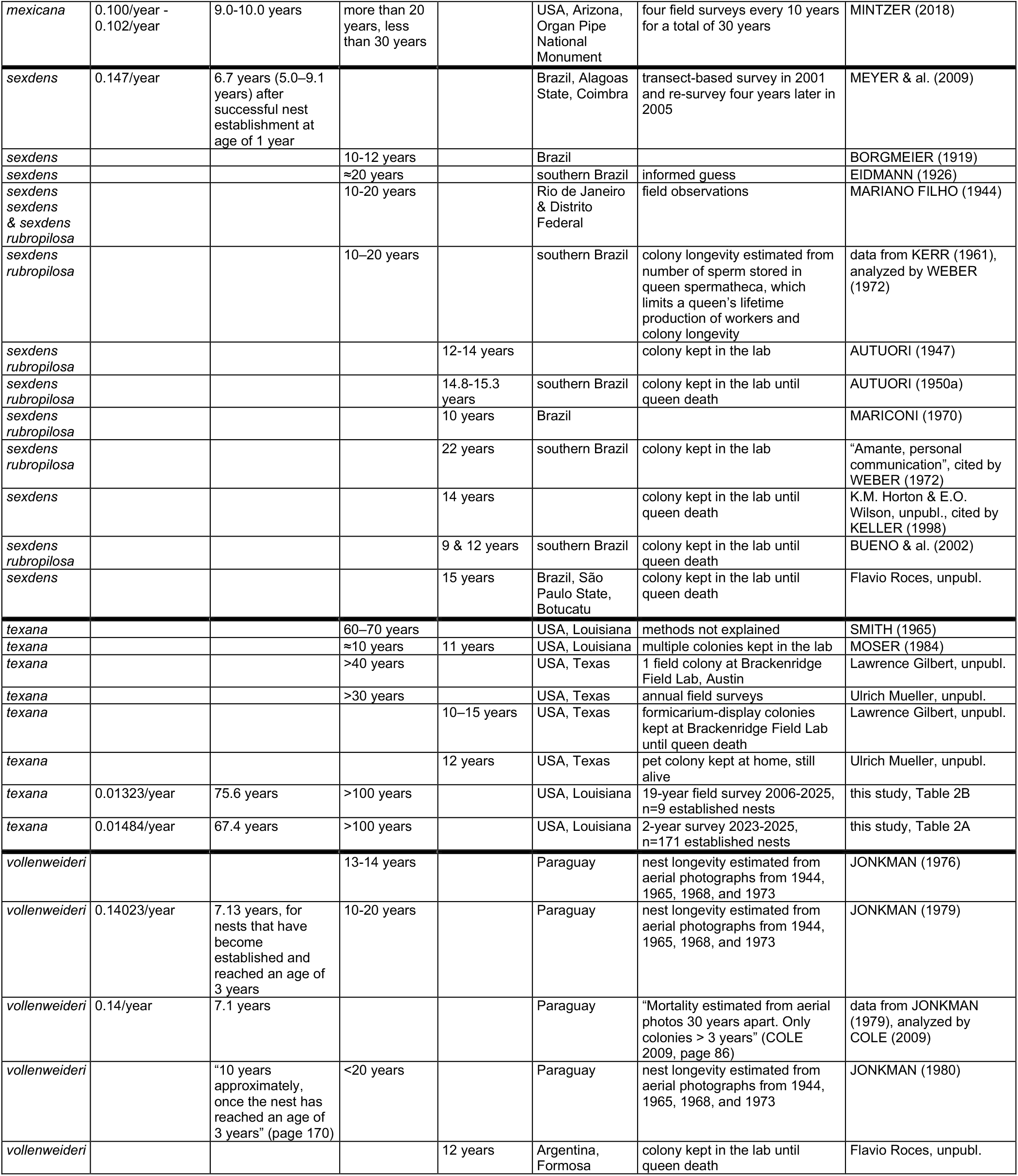
Annual *Atta* colony mortality after nest establishment in the field; average colony life expectancy after establishment in the field; maximum colony longevity in the field; and longevity of *Atta* queens in laboratory colonies. For studies that reported an annual mortality rate for established colonies (second column from left), or that reported information that allowed us to calculate the annual mortality rate, we estimate average colony life expectancy in the third column as the inverse of mortality rate (µ): average life expectancy = 1/µ. These life-expectancy estimates assume a constant colony mortality rate that is independent of colony age once a colony is established. All estimates of maximum queen longevity (fifth column) are based on observations of queens in laboratory colonies, not in field colonies. Table S2 in the digital supplementary materials includes footnotes to this table, including the details of our calculations of average colony life expectancies.

An unusual estimate of colony longevity was suggested in a little-known report by the myrmecologist Marion Smith for an *Atta texana* population at the range limit of the genus in north-west Louisiana, where Smith thought that single “colonies may occupy the same location for 60-70 years” (SMITH 1965, page 57). Smith was employed in 1936/1937 at the Southern Forest Experiment Station in western Louisiana (Provencal; now Kisatchie Ranger District) to develop strategies to eradicate *A. texana* (M.R. SMITH 1939, D.R. SMITH 1983), and his later report (SMITH 1965) did not explain his methods for deriving his estimate of 60-70 years colony longevity. Smith’s estimate of exceptional longevity in a range-limit population is surprising, because survivorship is often reduced at range limits due to occasional extreme environmental stresses that increase mortality, shorten lifespans, and maintain the range limit (GASTON 2009). Estimates of 50-year longevity for *A. texana* colonies were thought to be inflated by the *Atta* expert John Moser (MOSER 1984). In our 27-year studies of *A. texana* across Texas and Louisiana, we found striking differences in colony longevity between populations. We report here our colony-longevity estimates for *A. texana* in Louisiana, where it is easier to survey colonies because colony mortality is among the lowest that we observed for any population. We found that Smith’s estimate of colony longevity of 60-70 years (SMITH 1965) appears to be an underestimate, and that some *A. texana* colonies in Louisiana grow likely older than 100 years (Fig. 1).

**Fig. 1.**
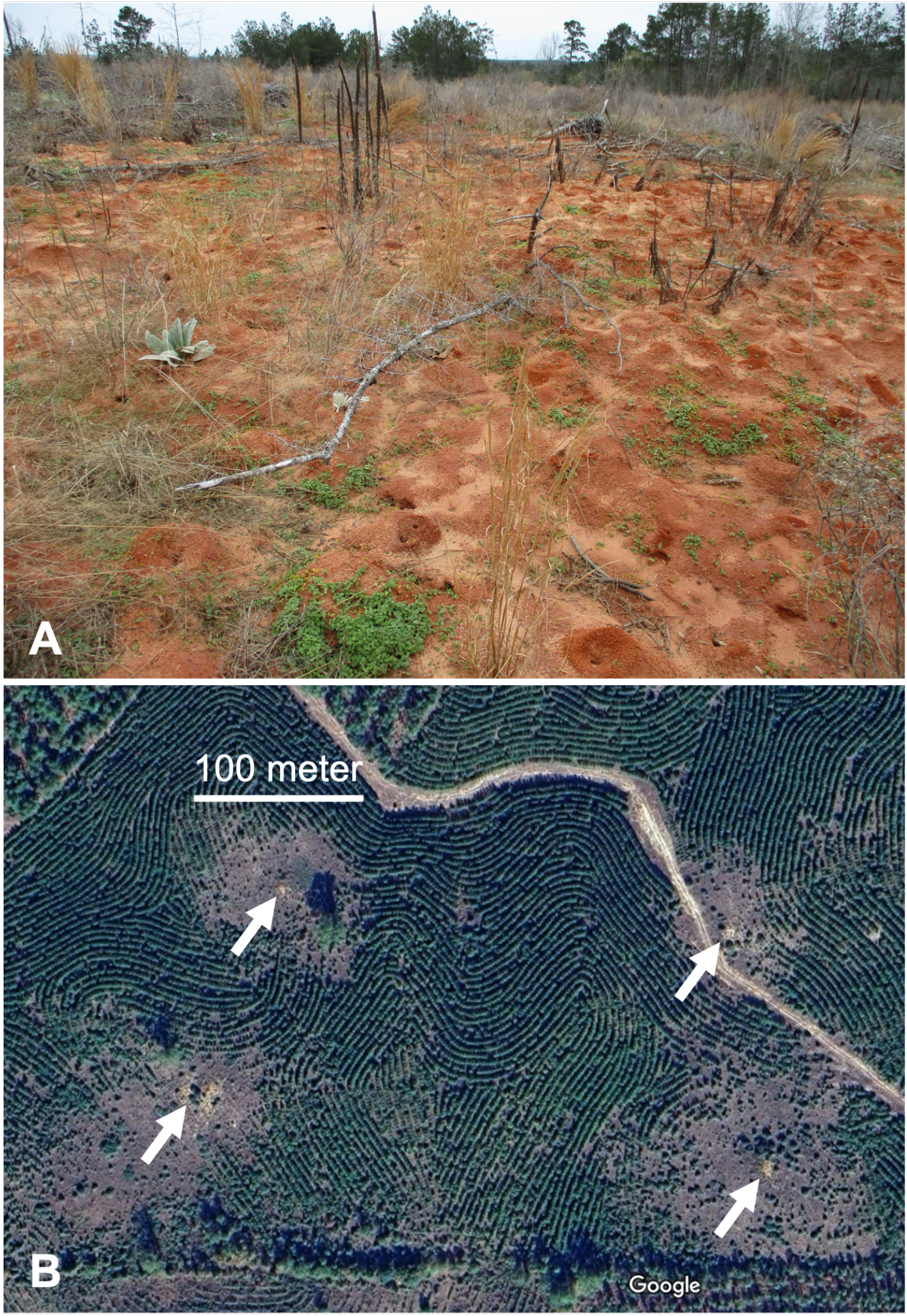
Nests of the leafcutter ant *Atta texana* in Louisiana. (A) *Atta texana* leafcutter mound measuring 32x29 m^2^ (size of two basketball courts combined, or size of a baseball infield diamond) in a pine plantation near Lucky, Louisiana. This single nest presumably has many queens that cooperate to produce the millions of workers of this giant colony. (B) Four large *A. texana* nests (arrows) in a young pine plantation, showing halos of open, sparsely-wooded areas within a 50-80 meter radius of each colony where newly-planted pines were destroyed by leafcutting foragers. Large *A. texana* nests are hyperdispersed because of between-colony competition, and their foraging territories are contiguous, so the actual foraging territories of the nests shown here reach well beyond their respective tree-less halos visible in this pine plantation. After their mating flights, thousands of newly-mated queens are attracted to such open areas to start their new nests, many of these queens are killed by workers of the resident colony, but some of these young queens may be adopted occasionally by a resident colony, thereby rejuvenating an old *A. texana* colony. Old colonies with large territories may recruit young queens especially well, such that large size attained in old age prolongs lifespan in *A. texana*.

## Materials and Methods

### Study species and strategy to find old, established colonies of *A. texana*

*Atta texana* is a fungus-growing ant species that can use diverse leaves (dicots, grass, pine needles) as substrate for the cultivation of fungus in underground garden chambers (WHEELER 1907, MOSER 1984, MEIRELLES & al. 2016, SCHOWALTER & RING 2017). Among the 15 recognized *Atta* species (BARRERA & al. 2022), *A. texana* is unusual because a single nest can have multiple queens (WALTER & al. 1938, MOSER 1962, MOSER 1963, ECHOLS 1966a, MOSER & LEWIS 1981, MINTZER & VINSON 1985, MINTZER 1987, U.G. Mueller, unpubl.); for example, seven queens were found in one large *A. texana* nest (MOSER 1963) and 16 queens (at least three of them fertilized) in a single garden of a 2-year-old nest (MOSER & LEWIS 1981). In contrast, newly-founded and mature nests of all other *Atta* species have been found, with very rare exceptions, with only a single queen (e.g., HUBER 1905a,b, EIDMANN 1936, STAHEL & GEIJSKES 1939, WEYRAUCH 1942, AUTUORI 1942, JACOBY 1944, WEBER 1956, WEBER 1972, FOWLER & al. 1984, FOWLER 1987, MINTZER 1991, MINTZER & al. 1991, NICHOLAS & VILELA 1995, SÁNCHEZ-PEÑA 2008, VIEIRA-NETO & al. 2016). Because workers of *A. texana* tolerate multiple queens in a nest, populations of *A. texana* likely exist as a mix of polygynous and monogynous nests, unlike the monogynous populations of all other *Atta* species. The proportion of polygynous versus monogynous nests is not known for any *A. texana* population. Polygynous nests of *A. texana* have been found in Louisiana, northern Texas, and central Texas (MOSER 1963, ECHOLS 1966a, MOSER & LEWIS 1981, U.G. Mueller, unpubl.), but so far not in southern populations of the *A. texana* range (U.G. Mueller, unpubl.). There exists no evidence that *A. texana* or any leafcutter ant species forms supercolonies (HELANTERÄ 2022). As in other well-studied ant genera with both monogynous and polygynous species (BOURKE & FRANKS 1995, HEINZE & FOITZIK 2009), polygynous colonies of *A. texana* are expected to exhibit different life-history traits compared to the monogynous colonies of other *Atta* species; for example, queen longevity in *A. texana* is predicted to be shorter than queen longevity in other *Atta* species (KELLER & GENOUD 1997, JEMIELITY & al. 2005, HEINZE & SCHREMPF 2008, JAIMES-NINO & OETTLER 2025).

In addition to polygyny, a second unique trait of *A. texana* is the capacity of different young nests to fuse sometimes in newly colonized habitat, thereby increasing growth rate of these merged, young colonies (ECHOLS 1966a), but experts dispute whether older colonies fuse, such that “any enlargement due to colony merging probably takes place only when colonies are young” (MOSER & LEWIS 1981, page 256). Third, the Louisiana population of *A. texana* is unique because of its recent origin and low genetic diversity (SMITH & al. 2019). After the end of the last glaciation ≈11,000 years ago, *A. texana* expanded across Texas and then into Louisiana (MUELLER & al. 2011a,b, SMITH & al. 2019), and serial founder effects during this recent range expansion can explain the observed low genetic diversity in Louisiana (SMITH & al. 2019). Low genetic diversity, in turn, may facilitate the merging of young *A. texana* colonies because they are genetically similar, resulting in inefficiency of genetically-based nestmate recognition such that young colonies sometimes fail to exclude non-nestmates and fuse nests (ECHOLS 1966a).

*Atta texana* is an edge-species (CAHAL 1993, SCHOWALTER & RING 2017) that is very rare in dense closed-canopy forest, but is most frequently found in semi-open forest or in forest gaps. Some of these gaps are created and maintained by the leaf-cutting activities of the ants (Fig. 1B). The central nest mound, where fungus gardens are concentrated underground (hereafter called “mound”), measures typically 7-15 meter diameter but can reach up to 35-40 meter diameter (Fig. 1A; Table S3) (SMITH 1939, CAHAL 1993, KULHAVY & al. 2001, SMITH 2001, U.G. Mueller, unpubl.). Such conspicuous mounds are easiest to find at forest edges along roads, gas pipelines, powerlines, or in Google Map satellite images (Fig. 1B). Only colonies with mounds measuring more than 7 meter diameter were selected for the below field surveys, to include only established colonies that had matured past the high-mortality stage of young, small colonies. It was easiest during field work to find larger mounds (>15 meter diameter) because they are linked via underground tunnels to many foraging exits in their correspondingly large foraging territories (Fig. 1B). The sample of surveyed nests is therefore biased towards colonies with large foraging territories and large prominent mounds (e.g., identifiable at Google Maps), and correspondingly likely biased also towards older colonies.

The present study uses commonly accepted definitions to distinguish between ant “nest” versus “colony”. An *A. texana* colony refers to the functional unit of millions of cooperating workers and reproductives, integrated by social organization, behavior, physiology (e.g., nutritional and hormonal interactions between ants), and species-specific nest architecture; whereas a nest is the physical structure constructed and inhabited by an *A. texana* colony, to serve as a defensible fortress buffered from ecological stresses, predators, parasites, and diseases (BOURKE & FRANKS 1995). *A. texana* is not known to be polydomous (i.e., an *A. texana* colony does not occupy more than one nest site at a time), except temporarily during the weeks or months when a colony migrates between two different locations within a colony’s territory, that is, when workers relocate brood and fungus gardens between pre-migration and post-migration nest sites. In Louisiana, migrations of *A. texana* colonies are very rare compared to populations in the southern range of *A. texana* (U.G. Mueller, unpubl.), and migrations of established colonies are typically triggered by extrinsic disturbances as defined by McGLYNN (2012). Only 6% of the surveyed *A. texana* colonies migrated during the present study, many of these migrations were back-and-forth migrations between two nest sites within a colony’s territory, and most of the colony migrations were triggered by disturbance by humans, such as logging of nearby forest or poisoning (see the accounts of all observed colony migrations described in detail in the digital supplementary materials). *A. texana* is the only leafcutter-ant species in Louisiana, and *A. texana* workers can be distinguished from those of other *Atta* species in the USA and Mexico by the matte, non-shiny appearance of head and gaster (SMITH 1963).

### Field surveys

Colony longevity was estimated in two surveys: (i) a 2-year survey of 195 established *A. texana* nests surveyed annually every January in 2023, 2024, and 2025 (Tables 2a and S3); and (ii) a 19-year survey of 9 established *A. texana* nests that were first geo-referenced in 2006 for genetic studies of the ant-cultivated fungi (MUELLER & al. 2011a,b) and that were re-surveyed again in 2019, 2020, and annually in 2022-2025 (Tables 2b, S1 and S3). The exact ages of these colonies at the start of each survey were not known, but because each nest was large (>7 meter diameter above-ground mound), each colony must have been at least 5-10 years old at the start of a survey (possibly much older, see below). Colonies included in the two surveys are therefore called here “established” colonies, defined as colonies that had survived the high-mortality phase of young colonies typical for the Type III survivorship of *Atta* (AUTUORI 1941, 1950b, MARICONI 1970, FOWLER & al. 1984, 1986, VIEIRA-NETO & VASCONCELOS 2010, FRÖHLE & ROCES 2012, MARTI & al. 2015) and typical for ants at large (COLE 2009, VERGARA-MARTíNEZ & al. 2024).

**Table 2.**
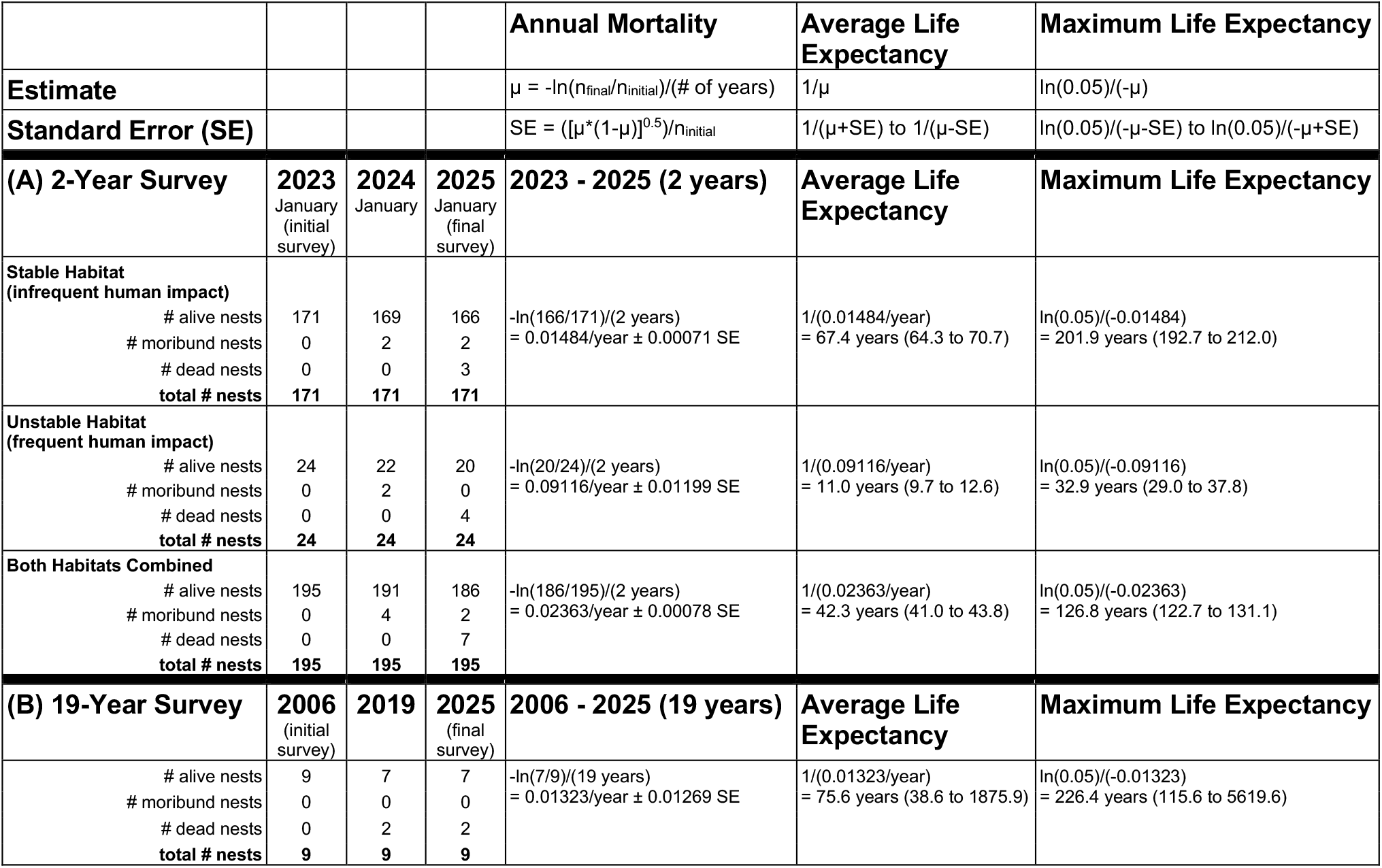
Mortality and life expectancy of *A. texana* nests in Louisiana estimated from a 2-year survey and a 19-year survey. (A) Mortality of 171 *A. texana* nests in stable habitat (infrequently impacted by humans) and 24 control nests in unstable habitat (sometimes impacted by humans; e.g., poisoning of *Atta* colonies), surveyed over two years every January in 2023-2025. Habitats were classified as stable or unstable before the 2024 re-survey (i.e., assignments to the two habitat types were pre-planned before scoring mortality). All seven nests scored dead in January 2025 were confirmed dead in April 2025. All four nests scored moribund in January 2024 were dead in January 2025. Two nests scored moribund in January 2025 were scored again as moribund three months later in April 2025. No nest scored moribund recovered and was found healthy in a later re-survey; therefore, once a nest is moribund, it appears to be effectively dead, and moribund nests are therefore pooled here with dead nests in the calculations of life expectancy. Life expectancy of nests is 6-times greater in stable, relatively undisturbed habitat compared to unstable habitat where nests are more frequently impacted by humans. (B) Mortality of 9 *A. texana* nests surveyed in Louisiana over 19 years between 2006-2025. These 9 nests were re-surveyed only six times at variable intervals between 2006-2025; see details of all censuses in the 19-year survey in Table S1 in the digital supplementary materials. SE = Standard Error.

The first sample of 9 nests of the 19-year survey includes all the *A. texana* nests known to the authors in 2006 in Louisiana. Four of these 9 nests were dug into in 2006 to reach the topmost fungus gardens (1.4-3.1 meter deep) to obtain live isolates of cultivated fungi for quantitative-genetic experiments (MUELLER & al. 2011a). Because of the large sizes of *A. texana* nests, digging a hole to access the topmost gardens and then refilling the hole should not have affected colony survivorship, but a minor impact on survivorship of these four dug-into colonies cannot be ruled out completely.

The 195 established nests of the 2-year survey were selected from 301 nests geo-referenced during annual field work from 2019 to 2023 in Louisiana. From these 301 geo-referenced nests, 106 nests were excluded because they were in locations with high likelihood of disturbance (e.g., yards of houses, old pine plantations approaching the time of logging), or nests where it would not be possible to search the surround within a 100-meter radius in all directions (e.g., a nest located next to a fenced-in property). Accessibility of a 100-meter radius around a nest was important for re-surveys to distinguish between migration versus mortality as causes of any colony disappearance. Both a dead mound and a mound abandoned by a migrating colony can look very similar (absence of fresh excavate on mound), but an *A. texana* colony that migrates has active foraging exits with fresh excavate at regular intervals between the abandoned mound and the post-migration mound (sometimes a few active exits remain for years on the abandoned mound, such that pre-migration and post-migration mounds remain linked via underground tunnels), whereas a colony that dies has no active foraging exits nearby the uninhabited mound. Fortunately, colony migrations in Louisiana are rare, and the rate of colony disappearance is low (see Results), so the re-surveys in 2024 and 2025 required few careful searches to distinguish between migration and death. For most nests, re-surveys required simply revisiting a nest’s GPS location to verify that a colony was still alive.

Each colony included in the baseline survey (2023) of the 2-year survey had large healthy worker populations, indicated by recent accumulations of excavate on the mound (Fig. 1A). Each time a nest was re-surveyed, visible signs were recorded indicating that the nest was still inhabited by a live *A. texana* colony, such as many excavating workers observable on the mound. The dimensions of the contiguous area of excavated soil defining the central mound (above-ground mound dimensions except radiating tunnels) were recorded at each visit, following the methods of CAHAL (1993), to quantify whether a colony was increasing in size, stable, or perhaps moribund (indicated by marked decrease in mound size compared to a year earlier). Mound size was estimated blind (KARDISH & al. 2015) with respect to any records from preceding years. The combined information allowed classification of each colony into (i) alive (ii) moribund, or (iii) dead. The digital supplementary materials includes field notes and measurements for all surveyed colonies (Table S3), as well as a more detailed account of all methods.

Of the 195 established nests included in the 2-year survey, and before the first re-survey in January 2024, 171 colonies were assigned to a Main Sample because these nests were judged to be in more stable and more protected habitat where disturbance by humans was expected to be infrequent (e.g., National Forest; younger pine plantations; semi-protected areas along gas pipelines or powerlines). For comparison, the remaining 24 nests were assigned to a Control Sample because these nests were judged to be in less stable and less protected habitat (e.g., in cemeteries, under roads) where disturbance by humans was expected to be more frequent (e.g., by poisoning or physical disturbance of leafcutter nests).

### Modeling of life expectancy and maximum lifespan

Annual mortality, life expectancy, and maximum life expectancy was estimated using the same approach to model these parameters in the primitive fungus-growing *ant Mycetosoritis hartmanni* (MUELLER & al. 2023). As in other studies of ant colony longevity (e.g., PAMILO 1991, MEYER & al. 2009, COLE 2009), this approach assumes that mortality is independent of age for established colonies (i.e., colonies that had survived the high-mortality stage of young colonies), and longevity can be estimated under a constant hazard model (i.e., colonies die off at a constant rate after establishment). The observed survivorship records from the 19-year survey (n=9 nests) and 2-year survey (n=171 main nests, n=24 control nests) were used to calculate estimates of the annualized mortality rate (μ, mortality per year) of nests. In the surveys, the number of surviving colonies declined every year due to mortality, so mortality across multiple years was estimated as μ = -ln(n_final_/n_initial_)/(# of years) [solving for μ from: n_final_ = n_initial_ * e^-μ*time^], where *n*_*final*_ is the number of nests alive at the end of a multi-year survey, *n*_*initial*_ is the number of nests alive at the beginning of the multi-year survey, and *# of years* is the total number of years elapsed between initial and final census. Assuming that mortality is constant with age for established colonies, this estimate of mortality μ was then used to calculate (i) an estimate of the expected lifespan (1/μ, the inverse of μ; i.e., the mathematical expectation of an exponential distribution); (ii) an estimate of the expected “maximum” lifespan as the age of the colonies when less than 5% of an initial cohort is expected to have survived [-μ^-1^ * ln(0.05), in years]; and (iii) the range of expected maximum lifespan derived from our estimate of mortality ± standard error of a binomial [1/(μ+SE_μ_) to 1/(μ-SE_μ_), in years], where SE_μ_ = ([μ * (1-μ)]^0.5^)/n_initial_ and *n*_*initial*_ is the total number of nests at the beginning of a survey. Constant annual colony survival likely approximates the true survival rate for much of the life of established colonies. In *A. texana*, colony survivorship does not appear to decline markedly in established nests after the death of the foundress queen(s) because large established nests adopt new queens regularly (see Results and Discussion).

## Results

### 19-Year Survey 2006–2025

Each of the 9 nests first surveyed in 2006 were established and large (>7 meter mound diameter), and these nests could be scored as unambiguously alive or dead in re-surveys (i.e., we observed no moribund nests in the 19-year survey; Table 2B; Table S1 in the digital supplementary materials). Because two of the 9 nests died during the 19-year survey from 2006*–*2025, the estimate of the annual mortality rate is - ln(7/9)/(19 years) = 0.01323/year. Assuming a constant mortality rate (i.e., mortality is independent of colony age), the average life expectancy of colonies can then be estimated as 1/(0.01323/year) = 75.6 years (Table 2B). Because queens of all *Atta* species, including *A. texana*, live for maximally 15-20 years (Table 1), such colony lifespans are only possible if colonies regularly adopt young queens to replace dying queens.

### 2-Year Survey 2023–2025

At the first-year mark in our 2-year survey, we observed in January 2024 no definitive cases of death among the 195 established nests, but we judged 4 nests to be moribund (2 moribund nests each among the 171 nests in stable habitat and among the 24 nests in unstable habitat, respectively; Table 2A). These 4 moribund nests were found dead one year later in our second re-survey in January 2025, suggesting that moribund colonies are more likely to die than to recover. In the second re-survey of our 2-year survey, we observed 2 moribund and 3 dead nests among the 171 nests in stable habitat, and 4 dead nests and no moribund nests among the 24 nests in unstable habitat (Table 2A). The two nests scored moribund in January 2025 were confirmed again as moribund three months later in April 2025, and all nests scored dead in January 2025 were confirmed dead in April 2025. No nest scored moribund recovered and was found alive and well in a later survey. Once a nest is moribund, therefore, the nest appears to be effectively dead, and we therefore pool moribund nests with dead nests in the below calculations.

For nests of *A. texana* in unstable habitat where nests are impacted more frequently by humans, we estimate a life expectancy of 11.0 years (Table 2A). This value is within the range of estimates reported for established nests of other *Atta* species (3.8*–*15.0 years life expectancy; Table 1). In contrast, our estimate of life expectancy of established *A. texana* nests in stable, relatively undisturbed habitat is about 6 times greater (67.4 years life expectancy) compared to unstable habitat (11.0 years life expectancy), because of lower annual mortality in stable habitat (1.484%) than unstable habitat (9.116%) (Table 2).

Our estimate of annual mortality rate of 1.484% in our 2-year survey (171 established nests in stable habitat) is higher than the mortality rate of 1.1323% estimated in our 19-year survey (9 established nests) (Table 2). Correspondingly, the estimate of average life expectancy is only 67.4 years for the 171 nests in our 2-year survey, lower than the 75.6 years life expectancy estimated for the 9 nests in our 19-year survey (Table 2). Assuming a constant hazard model of mortality (i.e., *A. texana* colonies die off at a constant rate once colonies have become established), the exponential decline of surviving colonies can be estimated by the survival function P(t) = e^−µt^, where µ is the mortality rate, and P(t) is the probability of *A. texana* colonies (or proportion of a cohort of colonies) surviving to at least an age of time t. Using this equation, the estimated mortality rate of µ=0.01323 predicts that 27% and 14% of the *A. texana* colonies should survive for, respectively, at least 100 or 150 years, well beyond the corresponding life expectancy of 75.6 years; and the estimated mortality rate of µ=0.01484 predicts that 23% and 11% of the *A. texana* colonies should survive, respectively, for at least 100 or 150 years, likewise well beyond the corresponding life expectancy of 67.4 years. Under a constant mortality rate of μ=0.01323, the maximum lifespan of colonies (lifespan attained by 5% of colonies in a population) can be estimated as -μ^-1^ * ln(0.05) = 226 years; under a constant mortality rate of μ=0.01484, the maximum lifespan of colonies can be estimated as -μ^-1^ * ln(0.05) = 202 years (Table 2A). These maximum colony-lifespan projections seem unrealistically high, because these projections assume that (i) the success of each colony in adopting young queens to replace dying queens does not change with time (e.g., there could be years when unusual weather patterns during the mating season reduces the chance of recruiting replacement queens); and (ii) no major catastrophes affect *A. texana* populations across centuries (e.g., there could be rare hurricane flooding that drowns many *A. texana* colonies throughout Louisiana).

## Discussion

Fascinating natural-history details hide often dormant in little-known niche publications, such as those by myrmecologist Marion Smith who hypothesized half a century ago for the leafcutter ant *Atta texana* in Louisiana that “colonies may occupy the same location for 60-70 years” (SMITH 1965, page 57). We found that Smith’s estimate of colony age appears to be an underestimate, and that some *A. texana* colonies in Louisiana likely can reach an age older than 100 years (Fig. 1, Table 2). Because *A. texana* queens live for less than 20 years (Table 1), colony lifespans older than 25 years are only possible if colonies adopt young queens to replace dying queens.

### *Atta texana* nests at the northern range limit have greater longevity than *Atta* nests in the tropics

Our 19-year survey (9 *A. texana* nests) and our 2-year survey (171 nests) estimate annual mortality rate of established *A. texana* colonies in Louisiana at only 1-2% per year (Table 2), at least 5-times lower than the mortality rates of 10-25% estimated for tropical *Atta* species (Table 1). A low annual colony mortality of less than 3% is unusual among ants, but it is plausible for *A. texana* because (i) we know of specific *A. texana* nests in central Texas that are at least 30-40 years old (likely much older, see footnotes of Table S2), confirming that *A. texana* colonies can requeen when older queens senesce and die; and (ii) annual colony mortality rates of less than 1% are known for ant species with multiple queens (JAIMES-NINO & OETTLER 2025), such as *A. texana*.

Because of low mortality of established *A. texana* colonies in Louisiana, we estimate that average colony-life expectancy is 5-10 times longer than for established colonies of other *Atta* species in tropical habitat (Table 1). Many nests of *A. texana* in Louisiana survive for decades in stable, undisturbed habitat, but not because *A. texana* has unusually long-lived queens (the known lifespan of 10-15 years of *A. texana* queens is comparable to queen lifespans of other *Atta* species; Table 1). Instead, workers of *A. texana* tolerate multiple queens in a colony (polygyny; WALTER & al. 1938, MOSER 1962, 1963, ECHOLS 1966a, MOSER & LEWIS 1981, MINTZER & VINSON 1985, MINTZER 1987, U.G. Mueller, unpubl.), and mature colonies adopt young, newly-mated queens on occasion, thereby requeening colonies as old queens die. Queen adoption may also enhance colony growth as supernumerary queens are added to a colony, which helps explain the many giant mounds of the largest *A. texana* colonies that we know, where the size of a single mound can reach 1000-1200 m^2^, the area of three basketball courts combined.

In contrast, colonies of tropical *Atta* species do not requeen at all, or perhaps do so at very low frequencies under natural conditions (WEYRAUCH 1942, AUTUORI 1947, 1950a, WEBER 1972, FOWLER & al. 1984, FOWLER 1987, MINTZER 1991, MINTZER & al. 1991, BUENO & al. 2002). Only two successful attempts to requeen an orphaned *Atta* laboratory colony have been reported so far (HUBER 1905b, SOTELO & al. 2015), most requeening experiments with tropical *Atta* species failed (EIDMANN 1936, AUTUORI 1950a, WEBER 1972, BUENO & al. 2002). Different from tropical *Atta* species where queen death leads to the inevitable demise of the colony (AUTUORI 1950a, WEBER 1982, WEISS 1990), queen death in a mature *A. texana* colony does not necessarily terminate the colony; instead, colony longevity in *A. texana* is determined by continuity of recruitment of young queens.

### Colony longevity and recovery of *Atta texana* populations from decade-long pesticide application

*Atta texana* has been studied in Louisiana for more than 90 years (HUNTER 1912, JONES 1917, SNYDER 1937, WALTER & al. 1938, SMITH 1939), so why has its centenarian colony longevity remained undocumented so far? A likely explanation is that, from the 1930s to the turn of the century, research and pest management prioritized eradication of *A. texana* as a forestry pest, and nests were routinely poisoned (with carbon disulfide or with arsenicals; SNYDER 1937, SMITH 1939), fumigated (with methyl-bromide gas; JOHNSTON 1944, MANN 1968), or killed by Mirex bait that was applied by aircraft across the landscape from 1962-1978 to control fire ants (ECHOLS 1966b, WILLIAMS & al. 2001, BARNETT & al. 2011). Because of landscape-wide, aerial pesticide application, “large mounds of *A. texana* disappeared” (BARNETT & BLOMQUIST 2020). After the organochloride Mirex was banned in the USA in 1978 because it is a bio-accumulating pollutant harming diverse animals, and after methyl-bromide was phased out between 1990-2005 because it acts as an ozone-depleting gas in the atmosphere, populations of *A. texana* recovered. Still, “it has taken nearly 50 years for their mounds to again become familiar sights in western Louisiana” (BARNETT & BLOMQUIST 2020) and for *A. texana* to regain its “earlier niche in the forest landscape” (BARNETT & al. 2011, page 41). Our study of colony longevity in *A. texana* therefore was fortuitously timely, and possible only after a 50-year post-Mirex recovery phase that allowed *A. texana* colonies in undisturbed habitat to grow old and live out their natural lifespans that are comparable to human lifespans.

### Large colony size attained in old age prolongs colony lifespan in *Atta texana*

Our 2-year survey was intentionally biased to include as many of the largest colonies that we could find (Fig. 1A), including colonies that would be near the end of their life if maximum colony lifespan of *A. texana* were about 20 years as in other *Atta* species (Table 1). Contrary to this expectation, it appears that the largest, most established *A. texana* colonies have great survivorship and longevity, possibly because such colonies have large foraging territories with large clearings around their nests (Fig. 1B), and these well-established colonies are therefore most likely to recruit newly-mated queens that are attracted to forest edges or clearings to start their new nests there. In *A. texana* in Louisiana, old age of a colony therefore does not necessarily increase mortality as in typical lifespan-limiting processes where age drives senescence and mortality (old age → mortality); instead, well-rooted residency that is correlated to age of an *A. texana* colony increases the chance of rejuvenation via queen adoption, reduces or eliminates reproductive senescence of a colony, and consequently prolongs colony lifespan (well-established residency → rejuvenation → long life). With respect to longevity and aging, therefore, *A. texana* in Louisiana appears similar to some other ant species where the largest colonies have highest survivorship (e.g., *Pogonomyrmex badius*, TSCHINKEL 2017) and similar to corals (BYTHELL & al. 2018) and many tree species (THOMAS 2013, CANNON & al. 2022, OHSE & al. 2023) where large size confers longevity benefits.

A prediction of the hypothesis of recurrent rejuvenation is that *A. texana* nests in closed-canopy forest (reduced chance of recruiting young queens because newly mated queens avoid founding their nests in closed-canopy forest) should have shorter colony lifespans than nests in forest gaps or at forest edges where young queens are most likely to settle, thereby facilitating queen adoption and rejuvenation of old colonies. We are currently testing this prediction in a much larger survey of 1097 established *A. texana* colonies distributed across the entire range of *A. texana*.

### Comparison with other polygynous ant and polygynous termite species

Within the genus *Atta, A. texana* is unique because it is the only species with significant rates of polygyny. Phylogenetic mapping onto the leafcutter-ant phylogeny (BACCI & al. 2009, Barrera & al. 2022) suggests that polygyny of *A. texana* likely arose from monogynous ancestors with multiply-mated queens (VILLESEN & al. 1999), most likely less than 1 million years ago (Barrera & al. 2022) and presumably near the northern range limit of the genus at that time. Concurrent with the evolution of polygyny, *A. texana* evolved a different social structure and unique life-history traits compared to all other *Atta* species, which all retained the putatively ancestral condition of monogyny during their evolution and diversification. In a larger comparative context, *Atta texana* may exhibit life-history traits that are convergent with traits exhibited by other polygynous ant species that likewise arose from monogynous ancestors, such as species in the well-studied ant genera with a mix of monogynous and polygynous species (e.g., *Solenopsis, Formica, Pogonomyrmex*; BOURKE & FRANKS 1995, HEINZE & FOITZIK 2009). For example, queen longevity in polygynous *A. texana* populations is predicted to be shorter than in the monogynous populations of all other *Atta* species (KELLER & GENOUD 1997, JEMIELITY & al. 2005, HEINZE & SCHREMPF 2008, JAIMES-NINO & OETTLER 2025). The available information on queen longevity is insufficient to test this prediction because of low sample sizes for *Atta* (Table 1) and because all estimates on *Atta* queen longevity reported so far are based on laboratory observations of queens in artificial nests that do not allow *Atta* queens to reproduce at rates typical for colonies in the field.

Serial polygyny and queen turnover mediating extraordinary colony longevity of *A. texana* has parallels with similar such social organization in some termite species with colony longevities reaching several decades and possibly much longer (BIGNELL & al. 2011, VARGO 2019, CHOUVENC & al. 2022). In such termite species, mating between reproductive offspring can occur inside the natal nest, resulting in continual turnover of reproductives within an “extended family” that resides for decades in the same termite nest (VARGO 2019). In contrast, *A. texana* serial polygyny occurs most likely because of regular adoption of queens that mate in typical mating flights outside their natal nests, specifically when newly-mated young queens found their incipient nests within the territory of a large established *A. texana* colony, in proximity of the adopting colony. Adoption of daughter queens is likely in Louisiana, because each older colony commands a large territory (Fig. 1) and because virgin queens of *A. texana* mate very close to their natal nest (essentially over the territory of their natal colony if the territory is large; Fig. 1), then descend immediately after mating to excavate their incipient nests (MOSER 1967, U.G. Mueller unpubl.). Depending on whether colonies adopt daughter queens or unrelated queens, *A. texana* populations may consist of a mix of “extended-family” and “mixed-family” colonies (sensu VARGO 2019). To test these scenarios of queen adoption that sustain serial polygyny in *A. texana* colonies, it will be necessary to genotype annually multiple neighboring nests of a single population, ideally over several decades, similar to the three-decade surveys by WIERNASZ & COLE (2026) of hundreds of nests of the harvester ant *Pogonomyrmex occidentalis* and by SUNDARAM & al. (2021) of *P. barbatus*. Such a survey of *A. texana* will be a major challenge, because it will be difficult to find a large number of *A. texana* nests in a single location, in natural habitat, and protected for several decades from human disturbance (see the digital supplementary materials for more information on human impacts causing colony mortality in *A. texana*).

### Immortality at the range limit

Both extrinsic and intrinsic factors likely contribute to exceptional longevity of *A. texana* colonies in Louisiana. First, specialized *Atta*-raiding *Nomamyrmex* army-ant predators (so-called “tank ants”) are absent in Louisiana, whereas *Nomamyrmex* army ants overlap with *A. texana* in south Texas and compromise colony survivorship there (KUBIK & al. 2024), as in tropical *Atta* species (BORGMEIER 1955, MARICONI 1970, SWARTZ 1998, RAO 2000, SÁNCHEZ-PEÑA & MUELLER 2002, LaPOLLA & al. 2002, POWELL & CLARK 2004, SOUZA & MOURA 2008). Second, because the genetic diversity within *A. texana* populations decreases along the post-glacial dispersal axis of *A. texana* from south Texas to Louisiana, such that range-limit populations in Louisiana are genetically most homogeneous (SMITH & al. 2019), the likelihood of re-queening may be intrinsically higher in Louisiana because of inefficiency of genetically-based nestmate-recognition, leading to a unique social structure within *A. texana* colonies, more frequent queen adoption, and perpetuation of serial polygyny. Recurrent queen adoptions make colonies potentially immortal. If so, leafcutter ants found their holy grail of colony immortality at their range limit in Louisiana, somewhere in the vicinity of a town called, appropriately, Lucky (N32.256° W92.997°).

## Acknowledgements

We thank the late John Moser for his advice to study the giant *Atta* mounds near Lucky, Louisiana; landowners for access to *Atta* nests; iNaturalist for location records; Flavio Roces and Lawrence Gilbert for their unpublished observations; Flavio Roces and two anonymous reviewers for exceedingly helpful comments improving our manuscript; the Wheeler Stengl-Lost-Pines Endowment and the National Science Foundation (awards DEB-1911443 to U.G.M., DEB-2239483 to C.E.F.) for financial support; and Casey Stengl for early encouragement to study the natural history of *Atta texana*.

## Declaration on use of generative artificial intelligence tools

The authors declare that they did not utilize generative artificial intelligence tools in any part of the analyses or the composition of this manuscript.

## Author contributions

Conceptualization: U.G.M., C.E.F. Methodology: U.G.M., C.E.F. Probability theory and analysis: C.E.F. Field surveys: U.G.M. Writing, original draft: U.G.M. Writing, review, and editing:

C.E.F. Funding acquisition: U.G.M., C.E.F.

## Data and code availability

All raw data are in Table S3 in the digital supplementary materials. The analyses did not require code but used the probability equations listed in Table 2.

## References

Autuori, M. 1941: Contribuição para o conhecimento da saúva (Atta spp. – Hymenoptera-Formicidae). I – Evolução do saúveiro (Atta sexdens rubropilosa Forel, 1908). – Arquivos do Instituto Biológico 12: 197–228.

Autuori, M. 1942: Contribuição para o conhecimento da saúva (Atta spp. – Hymenoptera-Formicidae). III – Excavação de um saúveiro (Atta sexdens rubropilosa Forel, 1908). – Arquivos do Instituto Biológico 13: 137–148.

Autuori, M. 1947: Contribuição para o conhecimento da saúva (Atta spp. – Hymenoptera-Formicidae). IV – O saúveiro depois da 1.a revoada (Atta sexdens rubropilosa Forel, 1908). – Arquivos do Instituto Biológico 18: 39–70.

Autuori, M. 1950a: Longevidade de uma colônia de saúva (Atta sexdens rubropilosa Forel, 1908) em condições de laboratório. – Ciência e Cultura 2: 825–826.

Autuori, M. 1950b: Contribuição para o conhecimento da saúva (Atta spp. Hymenoptera-Formicidae) V. Número de formas aladas e redução dos sauveiros iniciais. – Arquivos do Instituto Biológico 19: 325–331.

Bacci Jr, M., Solomon, S.E., Mueller, U.G., Martins, V.G., Carvalho, A.O., Vieira, L.G., & Silva-Pinhati, A.C.O. 2009: Phylogeny of leafcutter ants in the genus Atta Fabricius (Formicidae: Attini) based on mitochondrial and nuclear DNA sequences. – Molecular Phylogenetics and Evolution 51: 427–437.

Barnett, J.P., Haywood, J.D., & Pearson, H.A. 2011: Louisiana’s Palustris Experimental Forest: 75 years of research that transformed the South. General Technical Report SRS-148. – United States Department of Agriculture Forest Service, Southern Research Station, Asheville, NC, USA, 64 pp.

Barnett, J., & Blomquist, S. 2020: Getting to bottom of ant world. Louisiana Forestry Association (24 November 2020) – < https://www.laforestry.com/single-post/getting-to-bottom-of-ant-world >, retrieved on 13 April 2026.

Barrera, C.A., Sosa-Calvo, J., Schultz, T.R., Rabeling, C., & Bacci Jr, M. 2022: Phylogenomic reconstruction reveals new insights into the evolution and biogeography of Atta leaf-cutting ants (Hymenoptera: Formicidae). – Systematic Entomology 47: 13–35.

Bass, M. 1997: The effects of leaf deprivation on leaf-cutting ants and their mutualistic fungus. – Ecological Entomology 22: 384–389.

Bignell, D.E., Roisin, Y., & Lo, N. 2011: Biology of termites: A modern synthesis. – Springer, Dordrecht, The Netherlands, 576 pp.

Borgmeier, T. 1919: Vôo nupcial. – Vozes de Petrópolis 13: 169–172.

Borgmeier, T. 1955: Die Wanderameisen der neotropischen Region (Hym. Formicidae). – Studia Entomologica 3: 1–717.

Bourke, A.F.G., & Franks, N. 1995: Social evolution in ants. – Princeton University Press, Princeton, NJ, USA, 529 pp.

Bruner, S.C., & ValdÉS Barry, F. 1949: Observaciones sobre la biología de la bibijagua (Hymenoptera: Formicidae). – Memorias de la Sociedad Cubana de Historia Natural 19: 135–154.

Bueno, O.C., Hebling, M.J., Schneider, M.O., & Pagnocca, F.C. 2002: Occurrence of winged forms of Atta sexdens rubropilosa Forel (Hymenoptera: Formicidae) in laboratory colonies. – Neotropical Entomology 31: 469–473.

Bythell, J.C., Brown, B.E., & Kirkwood, T.B.L. 2018: Do reef corals age? – Biological Reviews 93: 1192–1202.

Cahal, R.R. 1993: The Texas leaf-cutting ant, Atta texana (Buckley) (Hymenoptera: Formicidae) and its effects on soil formation on sandy sites in East Texas. Master’s Thesis. – Stephen F. Austin State University, Nacogdoches, TX, USA, 73 pp.

Cole, B.J. 2009: The ecological setting of social evolution: The demography of ant populations. In: Gadau, J. & Fewell, J. (Eds.): Organization of insect societies: From genome to sociocomplexity. – Harvard University Press, Cambridge, MA, USA, pp. 74–104.

Cannon, C.H., Piovesan, G. & MunnÉ-Bosch, S. 2022: Old and ancient trees are life history lottery winners and vital evolutionary resources for long-term adaptive capacity. – Nature Plants 8: 136–145.

Chouvenc, T., Ban, P.M., & Su, N.Y. 2022: Life and death of termite colonies, a decades-long age demography perspective. – Frontiers in Ecology and Evolution 10: art. 911042.

Da Silva, J. 2022: The extension of foundress life span and the evolution of eusociality in the Hymenoptera. – American Naturalist 199: E140–E155.

Den Boer, S.P., Baer, B., Dreier, S., Aron, S., Nash, D.R., & Boomsma, J.J. 2009: Prudent sperm use by leaf-cutter ant queens. – Proceedings of the Royal Society B 276: 3945–3953.

Echols, H.W. 1966a: Compatibility of separate nests of Texas leaf-cutting ants. – Journal of Economic Entomology 59: 1299–1300.

Echols, H.W. 1966b: Texas leaf-cutting ant controlled with pelleted Mirex bait. – Journal of Economic Entomology 59: 628–631.

Eidmann, H. 1926: Brasilianische Skizzen. Die Blattscheiderameise und andere Forstschädlinge. – Forstwissenschaftliches Centralblatt 48: 593–613.

Eidmann, H. 1936: Das Atta-Problem: Untersuchungen über die Biologie und wirtschaftliche Bedeutung der Blattschneiderameise Atta sexdens L. – Die Naturwissenschaften 24: 257–266.

Fjerdingstad, E.J., & Boomsma, J.J. 1998: Multiple mating increases the sperm stores of Atta colombica leafcutter ant queens. – Behavioral Ecology and Sociobiology 42: 257–261.

Fowler, H.G. 1984: Population dynamics of the leaf-cutting ant, Atta capiguara, in Paraguay. – Ciência e Cultura 36: 628–632.

Fowler, H.G. 1987: Colonization patterns of the leaf cutting ant, Atta bisphaerica Forel: Evidence for population regulation. – Journal of Applied Entomology 104: 102–105.

Fowler, H.G., Pereira Da Silva, V., Forti, L.C., & Saes, N.B. 1986: Population dynamics of leaf-cutting ants: A brief review. In: Lofgren, C.S., & Vander Meer, R.K. (Eds.): Fire ants and leaf-cutting ants: Biology and management. – Westview Press, Boulder, CO, USA, pp. 123–145.

Fowler, H.G., Robinson, S.W., & Diehl, J. 1984: Effect of mature colony density on colonization and initial colony survivorship in Atta capiguara, a leaf-cutting ant. – Biotropica 16: 51–54.

FrÖHle, K., & Roces, F. 2012: The determination of nest depth in founding queens of leaf-cutting ants (Atta vollenweideri): idiothetic and temporal control. – Journal of Experimental Biology 215: 1642–1650.

Gaston, K.J. 2009: Geographic range limits: Achieving synthesis. – Proceedings of the Royal Society B: Biological Sciences 276: 1395–1406.

Heinze, J. & Foitzik, S. 2009: The evolution of queen numbers in ants: From one to many and back. In: Gadau, J. & Fewell, J. (Eds.): Organization of insect societies: From genome to sociocomplexity. – Harvard University Press, Cambridge, MA, USA, pp. 26–50.

Heinze, J., & Schrempf, A. 2008: Aging and reproduction in social insects: A mini-review. – Gerontology 54: 160–167.

HelanterÄ, H. 2022: Supercolonies of ants (Hymenoptera: Formicidae): Ecological patterns, behavioural processes and their implications for social evolution. – Myrmecological News 32: 1–22.

Huber, J. 1905a: Über die Koloniegründung bei Atta sexdens, (Erster Teil). – Biologisches Centralblatt 25: 606–619.

Huber, J. 1905b: Über die Koloniegründung bei Atta sexdens, (Schluss). – Biologisches Centralblatt 25: 625–635.

Hunter, W.D. 1912: Two destructive Texas ants. Reprinted 1922. Circular 148. – United States Department of Agriculture Bureau of Entomology, Washington, DC, USA, 7 pp.

Jacoby, M. 1944: Possibilidade da existência de duas rainhas em um único sauveiro. – Boletim da Sociedade Brasileira de Agronomia 7: 41–44.

Jaimes-Nino, L.M., & Oettler, J. 2025: The pace and shape of ant ageing. – Biological Reviews 100: 2071–2083.

Jemielity, S., Chapuisat, M., Parker, J.D., & Keller, L. 2005: Long live the queen: Studying aging in social insects. – Age 27: 241–248.

Johnston, H.R. 1944: Control of the Texas leaf-cutting ant with methyl bromide. – Journal of Forestry 42: 130–132.

Jones, T.H. 1917: Occurrence of a fungus-growing ant in Louisiana. – Journal of Economic Entomology 10: 561.

Jonkman, J.C.M. 1976: Biology and ecology of the leaf-cutting ant Atta vollenweideri, Forel 1893. – Zeitschrift für Angewandte Entomologie 81: 140–148.

Jonkman, J.C.M. 1979: Population dynamics of leaf-cutting ant nests in a Paraguayan pasture. – Zeitschrift für Angewandte Entomologie 87: 281–293.

Jonkman, J.C.M. 1980: The external and internal structure and growth of nests of the leaf-cutting ant Atta vollenweideri Forel, 1893 (Hym.: Formicidae) Part I. – Zeitschrift für Angewandte Entomologie 89: 158–173.

Kardish, M.R., Mueller, U.G., Amador-Vargas, S., Dietrich, E.I., Ma, R., Barrett, B., FANG, & C.C. 2015: Blind trust in unblinded observation in ecology, evolution, and behavior. – Frontiers in Ecology and Evolution 3: art. 51.

Keller, L. 1998: Queen lifespan and colony characteristics in ants and termites. – Insectes Sociaux 45: 235–246.

Keller, L., & Genoud, M. 1997: Extraordinary lifespans in ants: a test of evolutionary theories of ageing. – Nature 389: 958–960.

Kerr, W.E. 1961: Ascalamento de rainhas com vários machos em duas espécies da tribu Attini (Hymenoptera, Formicoidea). – Revista Brasileira de Biologia 21: 45–48.

Kubik, T.D., Mueller, U.G., Gibson, S., & Golightly, P. 2024: Fight or flight: defense strategies of fungus-farming ants (Attina) against the ant predators Nomamyrmex and Neivamyrmex. – Southwestern Entomologist 49: 1089–1104.

Kulhavy, D.L., Smith, L.A., & Ross, W.G. 2001: Impact of the Texas leaf-cutting ant (Atta texana (Buckley)) (Order Hymenoptera, Family Formicidae) on a forested landscape. In: Liebhold, A.M., Mcmanus, M.L., Otvos, I.S., & Fosbroke, S.L.C. (Eds): Proceedings: Integrated management and dynamics of forest defoliating insects; 1999 August 15–19; Victoria, BC. Gen. Tech. Rep. NE-277.

– United States Department of Agriculture Forest Service, Northeastern Research Station, Newtown Square, PA, USA, pp. 85–90.

Lapolla, J.S., Mueller, U.G., Seid, M., & Cover, S.P. 2002: Predation by the army ant Neivamyrmex rugulosus on the fungus-growing ant Trachymyrmex arizonensis. – Insectes Sociaux 49: 251–256.

Mann, W.F. 1968: Reinfestation of forest sites by the Texas leaf-cutting ant. – Journal of Economic Entomology 61: 1460–1461.

MARIANO Filho J. 1944: Contribuição ao conhecimento da biologia de algumas espécies do género Atta. – Boletim do Ministério da Agricultura 33: 19–29.

Mariconi, F.A.M. 1970: As saúvas. – Editora Agronômica Ceres, São Paulo, SP, Brazil, 167 pp.

Marti, H.E., Carlson, A.L., Brown, B.V., & Mueller, U.G. 2015: Foundress queen mortality and early colony growth of the leafcutter ant, Atta texana (Formicidae, Hymenoptera). – Insectes Sociaux 62: 357–363.

Mcglynn, T.P. 2012: The ecology of nest movement in social insects. – Annual Review of Entomology 57: 291–308.

Meirelles, L.A., Mcfrederick, Q.S., Rodrigues, A., Mantovani, J.D., De Melo Rodovalho, C., Ferreira, H., Bacci Jr, M., & Mueller, U.G. 2016: Bacterial microbiomes from vertically transmitted fungal inocula of the leaf-cutting ant Atta texana. – Environmental Microbiology Reports 8: 630–640.

Meyer, S.T., Leal, I.R., & Wirth, R. 2009: Persisting hyper-abundance of leaf-cutting ants (Atta spp.) at the edge of an old Atlantic forest fragment. – Biotropica 41: 711–716.

Mintzer, A. 1987: Primary polygyny in the ant Atta texana: number and weight of females and colony foundation success in the laboratory. – Insectes Sociaux 34: 108–117.

Mintzer, A. 2018: Changes over 30 years in populations of the leafcutter ant Atta mexicana at Organ Pipe Cactus National Monument. – Park Science 34: 32–42.

Mintzer, A. & Vinson, B. 1985: Cooperative colony foundation by females of the leafcutting ant Atta texana in the laboratory. – Journal of the New York Entomological Society 93: 1047–1051.

Mintzer, A.C. 1991: Colony foundation in leafcutting ants: the perils of polygyny in Atta laevigata (Hymenoptera: Formicidae). – Psyche 98: 1–5.

Mintzer, A., Quiroz-Robledo, L.N., & Deloya, C. 1991: Foundation of colonies of Atta Mexicana (F. Smith) (Hymenoptera: Formicidae) in the laboratory. – Folia Entomológica Mexicana 82: 133–138.

Moser, J.C. 1962: Probing the secrets of the town ant. – Forests & People 12: 12–13 & 40-41.

Moser, J.C. 1963: Contents and structure of Atta texana nest in summer. – Annals of the Entomological Society of America 56: 286–291.

Moser, J.C. 1967: Mating activities of Atta texana (Hymenoptera, Formicidae). – Insectes Sociaux 14: 295–312.

Moser, J.C. 1984: Town ant. In: Payne, T.L., Billings, R.F., Coulson, R.N., & Kulhavy, D.L. (Eds.): History, status, and future needs for entomology research in southern forests. Proceedings of the 10th Anniversary of the East Texas Forest Entomology Seminar, October 6–7, 1983.

Miscellaneous Publication MP-1553. – Texas Agricultural Experiment Station, Kurth Lake, TX, USA, pp. 47–52.

Moser, J.C., & Lewis, J.R. 1981: Multiple nest queens of Atta texana (Buckley 1860) (Hymenoptera: Formicidae). – Turrialba 31: 256–257.

Mueller, U.G., Himler, A.G., & Farrior, C.E. 2023: Life history, nest longevity, sex ratio, and nest architecture of the fungus-growing ant Mycetosoritis hartmanni (Formicidae: Attina). – Public Library of Science One 18: art. e0289146.

Mueller, U.G., Mikheyev, A.S., Hong, E., Sen, R., Warren, D.L., Solomon, S.E., Ishak, H.D., Cooper, M., Miller, J.L., Shaffer, K.A., & Juenger, T.E. 2011a: Evolution of cold-tolerant fungal symbionts permits winter fungiculture by leafcutter ants at the northern frontier of a tropical ant-fungus symbiosis. – Proceedings of the National Academy of Sciences USA 108: 4053–4056.

Mueller, U.G., Mikheyev, A.S., Solomon, S.E., & Cooper, M. 2011b: Frontier mutualism: coevolutionary patterns at the northern range limit of the leaf-cutter ant-fungus symbiosis. – Proceedings of the Royal Society B 278: 3050–3059.

Nicholas, J.T., & Vilela, E.F. 1995: Territorial mechanisms in post-nuptial flight gynes of the leaf-cutting ant Atta laevigata (F. Smith). – Anais da Sociedade Entomológica do Brasil 24: 389–400.

Ohse, B., Compagnoni, A., Farrior, C.E., Mcmahon, S.M., Salguero-GóMez, R., RüGer, N., & Knight, T.M. 2023: Demographic synthesis for global tree species conservation. – Trends in Ecology and Evolution 38: 579–590.

Pamilo, P. 1991: Life span of queens in the ant Formica exsecta. – Insectes Sociaux 38: 111–119.

Perfecto, I. & Vandermeer, J. 1993: Distribution and turnover rate of a population of Atta cephalotes in a tropical rain forest in Costa Rica. – Biotropica 25: 316–321.

Powell, S., & Clark, E. 2004: Combat between large derived societies: a subterranean army ant established as a predator of mature leaf-cutting ant colonies. – Insectes Sociaux 51: 342–351.

Rao, M. 2000: Variation in leaf-cutter ant (Atta sp.) densities in forest isolates: The potential role of predation. – Journal of Tropical Ecology 16: 209–225.

Salzemann, A., & Jaffe, K. 1990: Territorial ecology of the leaf-cutting ant, Atta laevigata. In: R.K. Vander Meer, K., Jaffe, K., Cedeno A. (Eds.): Applied myrmecology: A world perspective. – Westview Press, Boulder, CO, USA, pp. 345–354.

SÁNchez-PeÑA, S.R. 2008: Observations on aggression in leaf-cutting ant females, Atta Mexicana (Hymenoptera: Formicidae) in Mexico. – Entomologial News 119: 541–544.

SÁNchez-PeÑA, S.R., & Mueller, U.G. 2002: A nocturnal raid of Nomamyrmex army ants on Atta leaf-cutting ants in Tamaulipas, Mexico. – Southwestern Entomologist 27: 221–223.

Schowalter, T.D., & Ring, D.R. 2017: Biology and management of the Texas leafcutting ant (Hymenoptera: Formicidae). – Journal of Integrated Pest Management 8: art. 16.

Smith, C.C., Weber, J.N., Mikheyev, A.S., Roces, F., Bollazzi, M., Kellner, K., Seal, J.N., & Mueller, U.G. 2019: Landscape genomics of an obligate mutualism: Concordant and discordant population structures between the leafcutter ant Atta texana and its two main fungal symbiont types. – Molecular Ecology 28: 2831–2845.

Smith, D.R. 1983: Dr. Marion Russell Smith 1894–1981, Obituary. – Proceedings of the Entomological Society of Washington 85: 628–630.

Smith, L.A. 2001: Impact of the Texas leaf-cutting ant (Atta texana Buckley) (Hymenoptera: Formicidae) on foliar nutrient concentrations in droughty sites in East Texas. Master’s Thesis. – Stephen F. Austin State University, Nacogdoches, Texas, USA, 110 pp.

Smith, M.R. 1939: The Texas leaf-cutting ant (Atta texana Buckley) and its control in the Kisatchie National Forest of Louisiana. Occasional Paper No. 84. – United States Department of Agriculture Forest Service, Southern Forest Experiment Station, New Orleans, LA, USA, 5 pp.

Smith, M.R. 1963: Notes on the leaf-cutting ants, Atta spp., of the United States and Mexico (Hymenoptera: Formicidae). – Proceedings of the Entomological Society of Washington 65: 299–302.

Smith, M.R. 1965: House-infesting ants of the eastern United States, their recognition, biology, and economic importance. – United States Department of Agriculture Technical Bulletin 1326: 1–105.

Snyder, T.E. 1937: Damage to the young pines by a leaf-cutting ant, Atta texana Buckley, in Louisiana. – Louisiana Conservation Review 6: 14–17.

Sotelo, G., Ortiz-Giraldo, D.S., RodríGuez, J., & Montoya-Lerma, J. 2015: Adoption of a surrogate artificial queen in a colony of Atta cephalotes (L.) (Hymenoptera: Formicidae) in Colombia. – Sociobiology 62: 613–614.

Souza, J.L.P., & Moura, C.A.R. 2008: Predation of ants and termites by army ants, Nomamyrmex esenbeckii (Formicidae: Ecitoninae) in the Brazilian Amazon. – Sociobiology 52: 399–402.

Sundaram, M., Steiner, E., & Gordon, D.M. 2022: Rainfall, neighbors, and foraging: The dynamics of a population of red harvester ant colonies 1988–2019. – Ecological Monographs 92: art. e1503.

Stahel, G., & Geijskes, D.C. 1939: Ueber den Bau der Nester von Atta cephalotes L. und Atta sexdens L. (Hym. Formicidae). – Revista de Entomologia 10: 27–78.

Swartz, M.B. 1998: Predation on an Atta cephalotes colony by an army ant, Nomamyrmex esenbeckii. – Biotropica 30: 682–684.

Thomas, H. 2013: Senescence, ageing and death of the whole plant. – New Phytologist 197: 696–711.

Tschinkel, W.R. 2017. Lifespan, age, size-specific mortality and dispersion of colonies of the Florida harvester ant, Pogonomyrmex badius. – Insectes Sociaux 64: 285–296.

Urich, F.W. 1923: Ants in relation to agriculture. – Proceedings of the Agricultural Society of Trinidad and Tobago. Society Paper No. 795: 227–235.

Vargo, E.L. 2019: Diversity of termite breeding systems. – Insects 10: art. 52.

Vergara-MartíNez, M.F., Otero-DíAz, B., & Fetter-Pruneda, I. 2024: Unlocking the secrets of reproductive longevity: The potential of social insects. – Reproduction 167: art. e240020.

Vieira-Neto, E.H.M. & Vasconcelos, H.L. 2010: Developmental changes in factors limiting colony survival and growth of the leaf-cutter ant Atta laevigata. – Ecography 33: 538–544.

Vieira-Neto, E.H.M., Vasconcelos, H.L, & Bruna, E.M. 2016: Roads increase population growth rates of a native leaf-cutter ant in Neotropical savannahs. – Journal of Applied Ecology 53: 983–992.

Villesen, P., Gertsch, P.J., Frydenberg, J., Mueller, U.G., & Boomsma, J.J. 1999: Evolutionary transition from single to multiple mating in fungus-growing ants. – Molecular Ecology 8: 1819–1825.

Walter, E.V., Seaton, L., & Mathewson, A.A. 1938: The Texas leaf-cutting ant and its control. Circular 494. – United States Department of Agriculture Bureau of Entomology and Plant Quarantine, Washington, DC, USA, 18 pp.

Weber, N.A. 1956: Symbiosis between fungus-growing ants and their fungus. – American Philosophical Society Yearbook 1955: 153–157.

Weber, N.A. 1972: Gardening ants, the Attines. – The American Philosophical Society, Philadelphia, PA, USA, 146 pp.

Weber, N.A. 1976: A ten-year laboratory colony of Atta cephalotes. – Annals of the Entomological Society of America 69: 825–829.

Weber, N.A. 1982: Fungus ants. In: HERMANN, H.R. (Ed.): Social insects. Vol. 4. – Academic Press, New York, NY, USA, pp. 255–363.

Weiss, B.A. 1990: Longitudinal observation of attine (Atta cephalotes isthmicola) ants (Hymenoptera; Formicidae). – Zoo Biology 9: 421–429.

Weyrauch, W. 1942: Las hormigas cortadoras de hojas del Valle de Chanchamayo. – Boletín de la Dirección de Agricultura y Ganadería 13: 204–259.

Wheeler, W.M. 1907: The fungus-growing ants of North America. – Bulletin of the American Museum of Natural History 23: 669–807.

Williams, D.F., Collins, H.L., & Oi, D.H. 2001: The red imported fire ant: An historical perspective of treatment programs and the development of chemical baits for control. – American Entomologist 47:146–159.

Wiernasz, D.C., & Cole, B.J. 2026: Ant queen longevity in the field. – Insectes Sociaux 73: 111–116.

Wirth, R., Herz, H., Ryel, R.J., Beyschlag, W., HöLldobler, B. 2003: Herbivory of leaf-cutting ants. A case study on Atta colombica in the tropical rainforest of Panama. – Springer, Berlin, Germany, 230 pp.

