## Supplementary Material for "Undying leafcutter-ant colonies: The oldest *Atta texana* leafcutter-ant colonies are estimated to grow older than 100 years"

**Digital Supplementary Materials**


| | |
|---|---|
| Page 2 | Table S1: Survival and mortality of 9 *A. texana* nests surveyed in Louisiana over 19 years from April 2006 to January 2025. |
| Pages 3-9 | Table S2: Estimates of life-history parameters of *Atta* species reported in the literature. Table S2 is identical to Table 1 in the main article, but Table S2 includes extensive footnotes detailing the methods of the cited *Atta* studies, as well as explanations of any calculations used to estimate life-history parameters. |
| Pages 10-24 | Detailed field methods and field notes. |
| Pages 25-38 | Table S3: Raw data of field observations of all *Atta texana* nests included in the study. |

**Table S1:** Mortality of 9 established *A. texana* nests surveyed in Louisiana over 19 years from April 2006 to January 2025. Empty cells indicate that a particular nest was not re-surveyed at that time. An asterisk indicates that this nest was dug into at that time to access one of the topmost fungus gardens to obtain a live fungal culture. Of the four nests dug into in 2006, two nests survived until 2025, but two of these four nests died between 2006 and 2019. It is possible that digging into these four nests in 2006 impacted their survival. However, because of the large sizes of *A. texana* nests (fungal gardens are 1.5–8 meter below ground in a nest; queens are thought to inhabit the deeper fungal chambers), digging a hole that affects less than 10% of the nest and that reaches only the topmost fungal gardens, then refilling the hole after garden collection, should not have affected colony survivorship, although a minor impact on survivorship cannot be ruled out completely. Nests that were included also in our separate 2-year survey in 2023-2025 (Table 2 in main article) are indicated by their respective nest-IDs (e.g., nest #0090; see the numbering of nests in Table S3). The original nest UGM060403-01 was not included in our 2-year survey in 2023-2025 because a new house was built nearby in 2021, but the nest has survived so far in a protected corner of that property. Because two of the 9 original nests died during the 19-year survey, the annual mortality rate can be estimated as -ln(7/9)/19 = 0.01323/year. Assuming a constant mortality rate that is independent of colony age for established *A. texana* colonies, the average life expectancy can then be estimated as 1/(0.01323/years) = 75.6 years (Table 2 in main article).

| **2006 Nest Code** | **2025 Nest Codes** | **2006 Apr** | **2006 Dec** | **2019 Jun** | **2020 Nov** | **2021 Mar** | **2022 Dec** | **2023 Jan** | **2024 Jan** | **2025 Jan** |
|---|---|---|---|---|---|---|---|---|---|---|
| UGM060403-01 | UGM250117-37 | alive | | alive | alive | | alive | | alive | alive |
| UGM060403-06 | n/a | alive* | | dead | dead | | | dead | dead | dead |
| UGM060402-03 | UGM250103-04, nest #0090 | alive | | | alive | alive | | alive | alive | alive |
| UGM060402-04 | UGM250103-11, nest #0095 | alive* | | | | | | alive | alive | alive |
| UGM060402-05 | UGM250104-02; nest #0097 | alive | | alive | alive | alive | | alive | alive | alive |
| UGM061223-02 | UGM250103-27; nest #0110 | | alive | | | alive | | alive | alive | alive |
| UGM061223-03 | UGM250103-28; nest #0111 | | alive* | | | alive | | alive | alive | alive |
| UGM060402-06 | n/a | alive | alive* | dead | dead | | | dead | dead | dead |
| UGM060402-07 | UGM250103-35; nest #0118 | alive | | alive | alive | alive | | alive | alive | alive |

**Table S2:** Annual *Atta* colony mortality after nest establishment in the field; average colony life expectancy after establishment in the field; maximum colony longevity in the field; and longevity of *Atta* queens in laboratory colonies. For studies that reported an annual mortality rate for established colonies (second column from left), or that reported information that allowed us to calculate the annual mortality rate, we estimate average colony life expectancy in the third column as the inverse of mortality rate (μ): average life expectancy = 1/μ. These life-expectancy estimates assume a constant colony mortality rate that is independent of colony age once a colony is established. All estimates of maximum queen longevity (fifth column) are based on observations of queens in laboratory colonies, not in field colonies. Table S2 is identical to Table 1 in the main article, but Table S2 includes footnotes below the table explaining the methods of each study.

| ***Atta* species** | **Annual colony mortality rate (μ) after nest establishment in field** | **Average colony life expectancy, after nest establishment in field** | **Maximum colony longevity in field** | **Maximum queen longevity observed in laboratory** | **Location** | **Methods** | **References** |
|---|---|---|---|---|---|---|---|
| *capiguara* | | ≈7.5 years | | | Paraguay | informed guess | FOWLER (1984) |
| *capiguara* | 0.15/year | 6.7 years | | | Paraguay | “Survival estimated from photos 10-year interval. Survival of large nests estimated from table as 0.2 for 10 years” (COLE 2009, page 87) | data from FOWLER (1984), analyzed by COLE (2009) |
| *capiguara* | | 10-12 years | | | Paraguay | informed guess | FOWLER & al. (1986) |
| *cephalotes* | 0.295/year | 3.4 years | ≈16 years | | Costa Rica, La Selva | survey of 74 colonies from July 1989 to July 1991 | PERFECTO & VANDERMEER (1993) |
| *cephalotes* | 0.26/year (range 0.14/year to 0.57/year between different sites) | 3.8 years | | | Costa Rica, La Selva | “Mortality rates 0.14 – 0.57, across sites combined, mortality measured over two-year interval; N = 74 colonies” (COLE 2009, page 86) | data from PERFECTO & VANDERMEER (1993), analyzed by COLE (2009) |
| *cephalotes* | 0.147/year | 6.7 years (5.0–9.1 years) after successful nest establishment at age of 1 year | | | Brazil, Alagoas State, Coimbra | transect-based survey in 2001 and re-survey four years later in 2005 | MEYER & al. (2009) |
| *cephalotes* | | 14–15 years | | | Guatemala | | Dix & Dix, unpubl., cited by FOWLER & al. (1986) |
| *cephalotes* | | | 10-20 years | | Trinidad & Tobago | field observations | URICH (1923) |
| *cephalotes* | | | | 5–10 years | | | WEBER (1972) |
| *cephalotes* | | | | 10.2 years | | colony kept in the lab until queen death | WEBER (1976) |
| *cephalotes* | | | | 10-12 years | | | WEBER (1982) |
| *cephalotes* | | | | 10.4 years | | colony kept in the lab until queen death | WEISS (1990) |
| *cephalotes* | | | | 18 years | Trinidad | colony kept in the lab, used at age of 18 years for lab experiments | BASS (1997) |
| *cephalotes* | | | | 13 years | | colony kept in the lab until queen death | D. Cherix, unpubl., cited by KELLER (1998) |
| *cephalotes* | | | | 13 years | Panamá, Gamboa | colony kept in the lab until queen death | Flavio Roces, unpubl. |
| *colombica* | 0.092/year | 10.9 years | | | Panamá, BCI | surveys of 51 colonies from May 1996 to October 1998 | WIRTH & al. (2003) |
| *colombica* | 0.095/year | 10.5 years | | | Panamá, BCI | “Survival from multiple censuses. N = 92 colonies, 2 years” (COLE 2009, page 87) | data from WIRTH & al. (2003), analyzed by COLE (2009) |
| *colombica* | | ≈ 10 years | | | Panamá, Gamboa | colony longevity estimated from number of sperm stored in queen spermatheca, which limits lifetime production of workers and colony longevity | FJERDINGSTAD & BOOMSMA (1998) |
| *colombica* | | 8.54 years | <18 years | | Panamá, Gamboa | colony longevity estimated from number of sperm stored in queen spermatheca, which limits a queen’s lifetime production of workers and colony longevity | DEN BOER & al. (2009) |
| *colombica* | | | | 17 years | Panamá, Gamboa | colony kept in the lab until queen death | Flavio Roces, unpubl. |
| *insularis* | | | | 6.25 years | Havana, Cuba | colony kept in the lab | BRUNER & VALDÉS BARRY (1949) |
| *laevigata* | | | “several decades” | | Venezuela | informed guess | SALZEMANN & JAFFE (1990) |
| *laevigata* | | | | 11 years | Brazil, São Paulo State, Botucatu | colony kept in the lab until queen death | Flavio Roces, unpubl. |

| | | | | | | | |
|---|---|---|---|---|---|---|---|
| *mexicana* | 0.100/year - 0.102/year | 9.0-10.0 years | more than 20 years, less than 30 years | | USA, Arizona, Organ Pipe National Monument | four field surveys every 10 years for a total of 30 years | MINTZER (2018) |
| *sexdens* | 0.147/year | 6.7 years (5.0–9.1 years) after successful nest establishment at age of 1 year | | | Brazil, Alagoas State, Coimbra | transect-based survey in 2001 and re-survey four years later in 2005 | MEYER & al. (2009) |
| *sexdens* | | | 10-12 years | | Brazil | | BORGMEIER (1919) |
| *sexdens* | | | ≈20 years | | southern Brazil | informed guess | EIDMANN (1926) |
| *sexdens sexdens* & *sexdens rubropilosa* | | | 10-20 years | | Rio de Janeiro & Distrito Federal | field observations | MARIANO FILHO (1944) |
| *sexdens rubropilosa* | | | 10–20 years | | southern Brazil | colony longevity estimated from number of sperm stored in queen spermatheca, which limits a queen's lifetime production of workers and colony longevity | data from KERR (1961), analyzed by WEBER (1972) |
| *sexdens rubropilosa* | | | | 12-14 years | | colony kept in the lab | AUTUORI (1947) |
| *sexdens rubropilosa* | | | | 14.8-15.3 years | southern Brazil | colony kept in the lab until queen death | AUTUORI (1950a) |
| *sexdens rubropilosa* | | | | 10 years | Brazil | | MARICONI (1970) |
| *sexdens rubropilosa* | | | | 22 years | southern Brazil | colony kept in the lab | "Amante, personal communication", cited by WEBER (1972) |
| *sexdens* | | | | 14 years | | colony kept in the lab until queen death | K.M. Horton & E.O. Wilson, unpubl., cited by KELLER (1998) |
| *sexdens rubropilosa* | | | | 9 & 12 years | southern Brazil | colony kept in the lab until queen death | BUENO & al. (2002) |
| *sexdens* | | | | 15 years | Brazil, São Paulo State, Botucatu | colony kept in the lab until queen death | Flavio Roces, unpubl. |
| *texana* | | | 60–70 years | | USA, Louisiana | methods not explained | SMITH (1965) |
| *texana* | | | ≈10 years | 11 years | USA, Louisiana | multiple colonies kept in the lab | MOSER (1984) |
| *texana* | | | >40 years | | USA, Texas | 1 field colony at Brackenridge Field Lab, Austin | Lawrence Gilbert, unpubl. |
| *texana* | | | >30 years | | USA, Texas | annual field surveys | Ulrich Mueller, unpubl. |
| *texana* | | | | 10–15 years | USA, Texas | formicarium-display colonies kept at Brackenridge Field Lab until queen death | Lawrence Gilbert, unpubl. |
| *texana* | | | | 12 years | USA, Texas | pet colony kept at home, still alive | Ulrich Mueller, unpubl. |
| *texana* | 0.01323/year | 75.6 years | >100 years | | USA, Louisiana | 19-year field survey 2006-2025, n=9 established nests | this study, Table 2B |
| *texana* | 0.01484/year | 67.4 years | >100 years | | USA, Louisiana | 2-year survey 2023-2025, n=171 established nests | this study, Table 2A |
| *vollenweideri* | | | 13-14 years | | Paraguay | nest longevity estimated from aerial photographs from 1944, 1965, 1968, and 1973 | JONKMAN (1976) |
| *vollenweideri* | 0.14023/year | 7.13 years, for nests that have become established and reached an age of 3 years | 10-20 years | | Paraguay | nest longevity estimated from aerial photographs from 1944, 1965, 1968, and 1973 | JONKMAN (1979) |
| *vollenweideri* | 0.14/year | 7.1 years | | | Paraguay | "Mortality estimated from aerial photos 30 years apart. Only colonies > 3 years" (COLE 2009, page 86) | data from JONKMAN (1979), analyzed by COLE (2009) |
| *vollenweideri* | | "10 years approximately, once the nest has reached an age of 3 years" (page 170) | <20 years | | Paraguay | nest longevity estimated from aerial photographs from 1944, 1965, 1968, and 1973 | JONKMAN (1980) |
| *vollenweideri* | | | | 12 years | Argentina, Formosa | colony kept in the lab until queen death | Flavio Roces, unpubl. |

### Table S2 Footnotes

**Autuori (1950a).** Autuori reports that a queen of *Atta sexdens rubropilosa* lived in a laboratory colony for approximately 15 years and 4 months, equivalent to 15.33 years. The queen was collected with her incipient nest (garden & workers, excavated from the ground) in June 1935, 18 months after her probable mating flight in December 1933, then kept in the laboratory until the death of the queen sometime between October 1948 (the month the queen was last seen alive) and April 1949 (the month the queen was found dead in the refuse pile of the nest). The laboratory nest was already deteriorating and the worker population and garden sizes were decreasing in October 1948. The actual estimate of queen longevity therefore ranges between 14.8 (queen last seen alive in October 1948) to 15.3 years (queen found dead in April 1949). The greater estimate of 15.3 years queen longevity has been cited consistently in the literature.

**Bass (1997).** Bass describes experiments with a colony of *Atta cephalotes* that had been kept in the laboratory for 18 years: "Experiments were carried out using a laboratory nest of *Atta cephalotes*, aged 18 years and originally from Trinidad. The nest had about 600 000 workers and 140 L of fungus garden built in 2.5-L plastic domes" (page 385).

**Cole (2009).** Table 4.3 on pages 86 & 87 of Cole (2009) summarizes gross annual mortality rates and extrapolated life expectancies for *Atta cephalotes*, *A. colombica*, *A. capiguara*, *A. vollenweideri*, and 10 additional ant species from 3 ant subfamilies.

**den Boer et al. (2009).** den Boer et al. improved the methods of Fjerdingstad & Boomsma (1998; see below) to (i) estimate the average number of sperm and between-queen variance of sperm stored in spermathecae of *Atta colombica* queens, and (ii) determine the number of sperm used by a queen to fertilize a single egg. The median estimate for the number of sperm used by a queen to fertilize a single egg was estimated to be two sperm per egg; "median sperm use itself was also higher in founding queens (7.17 ± 1.58 s.e.) than in established queens (2.73 ± 0.34)… For established queens, median sperm use increased with queen age" (page 3949). Using these estimates of sperm use, den Boer et al. then modeled sperm use by a queen under typical colony growth in the field, to predict the queen age when a queen exhausts her sperm stores and the resulting cessation of worker production leads to colony demise. "We used the leaf-cutter ant *Atta colombica* to investigate the dynamics of sperm use during egg fertilization. We show that queens are able to fertilize close to 100 per cent of the eggs and that the average sperm use per egg is very low, but increases with queen age. The robustness of stored sperm was found to decrease with years of storage, signifying that senescence affects sperm either directly or indirectly via the declining glandular secretions or deteriorating sperm-storage organs. We evaluate our findings with a heuristic model, which suggests that the average queen has sperm for almost 9 years of normal colony development" (page 3945). den Boer et al.'s "model assumes that the average *A. colombica* colony goes through a period of logistic growth during the first 5 years after establishment before achieving a size of two million workers, after which 3000 gynes are produced each year ... The initial number of sperm stored was drawn randomly from a normal distribution with a mean of 244 million and standard deviation of 83 million … , and the average worker lifespan was set at four months … , so that the maintenance of a worker force of two million would require the annual fertilization of six million eggs. Stored sperm viability was assumed to be 100 per cent throughout. Sperm use and remaining sperm supplies were calculated for each six-month period after colony founding, based on different combinations of the initial sperm use per egg and the observed change in this number with age, so that the expected age at which queens run out of sperm could be estimated. The model was run 10 000 times to determine whether the patterns of sperm use and storage that we observed were consistent with the model" page 3747). Figure 4 (page 3950) summarizes the model predictions of queen and colony longevity, based entirely on sperm depletion. Under the most conservative assumptions of sperm use to fertilize a single egg, Figure 4 shows that the maximum age that an *A. colombica* colony can attain in the field is 18 years. "The results of our heuristic model suggest that sperm limitation is likely to lead to an average queen and colony longevity of 8.54 years, meaning that sexuals will, on average, only be produced for 3–4 years and that 5 per cent of colonies will run out of sperm before producing any sexuals" (page 3951).

**Eidmann (1926).** Eidmann writes about *Atta* nests on page 601: "Bei alten Nestern (von vielleicht 20 Jahren oder mehr)…" [translation: "In old nests (of perhaps 20 years or older…)], indicating that Eidmann believes that *Atta* nests can reach an age of about 20 years. In his later publications, Eidmann never specifies exact or approximate estimates for ages of old *Atta* nests, but he refers more generally to old nests ("alt") and super-old nests ("uralt"; literally "ancient"), without giving approximate ages for old and ancient nests.

**Fjerdingstad & Boomsma (1998).** Using the same approach as Weber (1972, page 38), Fjerdingstad & Boomsma (1998) estimate the number of sperm stored in the spermatheca of a mated female *Atta colombica* to estimate the number of workers that a queen can potentially produce during her lifetime, and from that estimate of worker number then project the number of years that a queen can potentially remain fertile to produce workers to sustain her colony. "The average queen [of *A. colombica*] had stored $100 \times 10^6$ sperm (range: 7-234 million, SD = $56 \times 10^6$)" (page 258). "A conservative estimate is that a queen which lives and reproduces for 10 years would need at least 80 ± 270 million sperm" (page 260). "The available life history data therefore indicate that, in natural populations, the number of sperm stored by an *A. colombica* queen is likely to constrain her reproductive lifespan and hence output" (page 260).

**Fowler (1984).** Fowler studied population dynamics of *Atta capiguara* in eastern Paraguay by comparing nest densities in a population with densities observed 10 years earlier in that same population (nest densities were inferred from aerial photographs). "Execution by workers of … founding queens is much higher in higher density areas, which, in effect, gives a distinct advantage to queens initiating colonies in less dense habitats" (page 631). "… characteristics of the habitat such as vegetation and density of colonies may ultimately determine colony survivorship. Indeed, the survivorship curves given here may be interpreted as density-dependent control of survivorship, which is carried out by the death of the weaker smaller colonies. Because of the survey techniques employed during this study, it is impossible to determine exactly how long a colony might live, as there are presently no data linking chronological age with nest dimensions, such as employed by Jonkman (1979) in his study of the survivorship of *Atta vollenweideri*. However, based on this study (Jonkman 1979), an average of 7.5 years may also be appropriate [for *Atta capiguara*]" (page 631). That is, Fowler suggests here that the average life expectancy of "7.5 years" for established colonies of *A. vollenweideri* is "appropriate" also for *A. capiguara*. (Jonkman 1979, see below, actually inferred 7.13 years nest longevity for *A. vollenweideri*, not 7.5 years longevity as claimed by Fowler.) Fowler bases his estimate of 7.5 years colony life expectancy of *A. capiguara* on his understanding of the natural history of *A. capiguara*, not on hard data collected during his population-dynamic investigations in eastern Paraguay.

**Fowler et al. (1986).** Fowler writes: "Dix and Dix (unpublished) … estimated the average age to maturity was five to six years for *A, cephalotes* and that colonies probably survived until the age of 14 to 15 years. Fowler (1984) studied the population dynamics of *A. capiguara* and found a Type I (heavy initial mortality) survivorship. In this species, colonies are estimated to live 10 to 12 years once they pass through the initial phase" (page 128). Fowler et al. (1986) incorrectly label here survivorship curves with "heavy initial mortality" as Type I; this is actually Type III survivorship. Fowler et al. (1986) offer a disparaging assessment of Schade's (1973) suggestion that maximum lifespan of *Atta sexdens rubropilosa* may exceed 100 years, writing this biting comment: "Schade (1973) referred to colony life spans of *Atta sexdens rubropilosa* on the order of 30 to 140 years, but apparently these estimates are not based on data" (page 128).

**Keller (1998).** Table 1 on pages 237-240 in Keller (1998) summarizes estimates of colony lifespan and queen lifespan for 54 ant species, including *A. cephalotes* and *A. sexdens*. For *A. sexdens*, Keller (1998, page 239) cites an unpublished record of queen longevity of 14 years in a laboratory colony maintained by Kathy Horton and Edward O. Wilson at Harvard University. For *A. cephalotes*, Keller (1998, page 240) cites an unpublished record of queen longevity of 13 years in a colony maintained in the laboratory of Daniel Cherix at the University of Lausanne.

**Jonkman (1976).** Figure 2 on page 143 indicates that Jonkman estimated the oldest nests of *Atta vollenweideri* to be 13-14 years old. Jonkman graphed in Figure 2 the age distribution approximately as 4.5-6.5 years old (n=27 nests), 7.5-8.5 years old (n=19 nests), 9.5-10.5 years old (n=4 nests), and 11.5-14.0 years old (n= 7 nests), suggesting that the percentage of older nests (> 10 years old) is less than the percentage of younger nests (5-10 years old) in this population in Paraguay. The location of this study population is not specified in Jonkman (1976), so it is unclear (but likely) whether these data are from the same population as the data reported in Jonkman (1979).

**Jonkman (1979).** Table 2 (page 288) categorizes live nests observed at study site "km 41" into four age classes and lists number of nests by age class: 4.5-6.5 years old (n=23 nests), 7.5-8.5 years old (n=14 nests), 9.5-10.5 years old (n=3 nests), and 11.5-20.0 years old (n= 5 nests). "The calculation revealed that a 3 year old colony has an average additional life expectancy of 7.13 years. Roughly speaking, this means that, once clearly established, colonies will reach an average age of about 10 years" (page 290). The Appendix on page 293 explains the exact calculations of how Jonkman derived the estimate of average life expectancy for established nests. The methods assume a constant mortality rate: "The paucity of the data make the assumption necessary that the probability of a colony, surviving one more year, is constant (= p), regardless of the age of the colony, as long as it is at least 3 years of age" (page 293). Jonkman estimates on page 293 the annual death rate ("m"; i.e., mortality) as 0.14023/year, and the average life expectancy as 1/(0.14023/year) = 7.13 years. Jonkman concludes that "… a nest aged 3 can be expected to survive 7.13 more years on the average" (page 293).

**Jonkman (1980, Part I).** "None of the nests in age class 4 [= the oldest nests found in aerial photographs in Jonkman's 1973 survey] appeared in a photograph of the same area made in 1944, which indicates that these nests did not become older than 30 years. The vegetation growth patterns on several of the oldest living nests indicates that they are probably not older than 20 years" (page 167). In his earlier publication Jonkman estimated an average life expectancy for *A. vollenweideri* colonies of 7.13 years (see above, Jonkman 1979), but in his publication a year later (Jonkman 1980), he summarizes his earlier research by giving a somewhat older estimate of average life expectancy for A. vollenweideri colonies: "… the average life expectancy, which is estimated to be 10 years approximately, once the nests have reached the age of 3 years (Jonkman 1979a)" (page 170).

**Mariano Filho (1944).** Mariano Filho writes on page 25 that *Atta sexdens* mounds can be 10-20 years old, but Mariano Filho (page 23) also considers the possibility of queen replacement, and that *Atta* colonies may live indefinitely: "A favor da hipótese, de que as formigas saúvas podem substituir as rainhas mortas ou inválidas, pode-se invocar o fato de que os formigueiros não morrem ou se extinguem. … o formigueiro mais velho, aquêle que inicialmente ocupou o terreno e forneceu as tanajuras para as otras formações, continuará a vivir indefinidamente" (translation: In support of the hypothesis that leafcutter ants can replace dead or decrepit queens, one can cite the fact that the ant colonies do not die out or become extinct. … the oldest ant colony, the one that initially colonized an area and provided the queen ants for new colonies, will continue to live indefinitely).

**Mariconi (1970).** Mariconi writes regarding longevity of *Atta* nests in the lab: "A manutenção de içás vivas, em sauveiros criado em laboratório, por 5, 7, e até 10 anos não é tão raro" (page 34) (translation: "Keeping live queens, in laboratory nests, for 5, 7 and even 10 years is not so rare). Regarding queen death Mariconi writes "… após a morte da içá, o sauveiro também morre" (page 34) (translation: "… after the queen dies, the colony also dies"). Mariconi reports on page 35 lab experiments on *Atta sexdens rubropilosa* and *Atta bisphaerica* indicating that new queens can be introduced to small young laboratory nests ("… substituir as rainhas de pequenas colónias por prazo que variou de 7 a cérca de 60 dias; translation: "… replace the queens of small colonies for a period that varied from 7 to around 60 days"). Mariconi concludes on page 35 that "… o pensamento geral, inclusive o nosso, é de que morrendo a rainha, o sauveiro morre" (translation: "the general thought is, including our [opinion], that once the queen dies, the *Atta* colony dies").

**Meyer et al. (2009).** Meyer et al. calculate an identical estimate of colony life expectancy for *A. sexdens* and *A. cephalotes*, because observations from both species were pooled in calculations (page 714) of colony life expectancy for these two sympatric *Atta* species. Because of the pooling of observations from two *Atta* species surveyed concurrently at the same site (forest in Coimbra, State of Alagoas, Brazil), the reported average colony life expectancy of 6.7 years (reported range of 5.0 - 9.1 years colony life expectancy) is the average across both species, and differences in colony life expectation between these two species cannot be inferred from the reported data. As in Perfecto and Vandermeer (1993), Meyer et al. surveyed transects along preexisting foot trails (total of 16 transects along forest edges and through interior forest). Transects were first surveyed between October 2001 and May 2002, and again between August and November 2005, a time span of approximately four years. "Of 36 [leafcutter ant] colonies recorded in 2001, 20 survived until 2005, 16 colonies died, and 19 new colonies were established" (page 713). Because birth rate is similar to death rate, the population appeared stable across the four survey years. Because 20 of 36 initial colonies survived until the 4-year re-survey, we calculate, using the equation in Table 2 (main article), an estimate of the annual colony mortality rate as $\mu = -\ln(20/36)/(4) = 0.14695$/year, and our estimate of colony life expectancy if therefore 1/(0.14695/year) = 6.8 years. Meyer et al. calculate colony life expectancy using a somewhat different method; they calculate colony life expectancy as the inverse of turnover rate ("calculated as the sum of new and dead colonies in 2005 divided by the sum of total colonies in 2001 and 2005", page 713), and Meyer et al. calculate turnover rates separately for transects along forest edges versus transects in interior forest. Meyer et al.'s estimate of colony life expectancy of "6.7 years (range 5.0 - 9.1 years)" (page 714), derived from turnover rate, is nearly identical to our estimate of 6.8 years colony life expectancy derived from annual colony mortality rate. Interestingly, "roughly half of all colonies were replaced by new ones during the 4-yr study period" (page 714), similar to the turnover rate of *Atta colombica* nests observed by Wirth et al. (2003) on Barro Colorado Island (BCI) in Panamá. Meyer et al. did not observe any colony migrations during the 4-year study (in contrast, 25% of the *Atta* colonies were observed to relocate annually on BCI Panamá). Meyer et al. assumed in their study that *Atta* colonies becomes established at an age of 1-year old, whereas most other studies for which information is summarized in Table S1 assume that *Atta* colonies become established sometime between a colony age of 2-4 years (i.e., other studies exclude the younger, establishing 1-year-old colonies that are not yet fully established). Because Meyer et al. include younger, establishing colonies that likely experience greater mortality than established colonies, their reported estimate of annual nest mortality likely overestimates the mortality rate of established colonies, and the colony life expectancy for established colonies is likely greater than the "6.7 years (range 5 - 9.1 years)" reported by Meyer et al.

**Mintzer (2018).** Mintzer writes that he geo-referenced 35 live *Atta mexicana* colonies in a 1985 survey at Organ Pipe National Monument (Arizona, USA), added 2 more large colonies in 1987 to this sample (for a total of 37 colonies), then re-surveyed these 37 colonies every 10 years over a 30-year time period. For the 20-year survey conducted in winter 2005/2006, Mintzer (2018) reports that "only four or five colonies first located in 1985 were still alive in 2005–2006, indicating that typical field colony life span is less than 20 years" (page 37). None of the original 37 colonies were alive in the 30-year survey conducted in winter 2015/2016 ("none has survived over the complete 30-year survey period from 1985 to 2016"; page 37), indicating that the maximum longevity of *Atta mexicana* colonies at Organ Pipe National Monument is greater than 20 years, but less than 30 years. In the 2015/2016 survey, "the great majority of colonies were younger than 10 years, and most were 1–2 or 7–10 years old" (page 37); colonies older than 10 years represented therefore a small minority in the 2015/2016 survey.

Because Mintzer writes that "only four or five colonies first located in 1985 were still alive" (page 37) after 20 years, we calculate here the colony mortality rate and average life expectancies separately for the case of 5 colonies surviving (32 of the initial 37 colonies died within 20 years) and the case of 4 colonies surviving (33 of the initial 37 colonies died within 20 years); and we include in our calculations the two large colonies added to the sample in 1987 (i.e., we assume that, because the two colonies added to the sample in 1987 were large, they must have been alive already two years earlier in 1985). We use the equations from our Table 2 to calculate estimates of annual mortality and average life expectancy:

***Assuming that 5 colonies of the initial 37 colonies survived for 20 years*:** Five surviving colonies after 20 years out of 37 initial colonies indicates an estimate of an annual mortality rate of $\mu = -\ln(n_{final}/n_{initial})/(\#\text{ of years}) = -\ln(5/37)/(20) = 0.10007$. Assuming a constant mortality rate, the estimate of the average life expectancy is $1/\mu = 1/(0.10007/\text{year}) = 10.0$ years; and the estimate of maximum colony longevity is $\ln(0.05)/(-\mu) = \ln(0.05)/(-0.10007) = 29.9$ years (5% of the colonies in this population may reach this age, assuming the foundress queen survives that long, or colonies can requeen after death of the foundress queen).

***Assuming that 4 colonies of the initial 37 colonies survived for 20 years*:** Four surviving colonies after 20 years out of 37 initial colonies indicates an estimate of an annual mortality rate of $\mu = -\ln(n_{final}/n_{initial})/(\#\text{ of years}) = -\ln(4/37)/(20) = 0.11123$. Assuming a constant mortality rate, the estimate of the average life expectancy is $1/\mu = 1/(0.11123/\text{year}) = 9.0$ years; and the estimate of maximum colony longevity is $\ln(0.05)/(-\mu) = \ln(0.05)/(-0.11123) = 26.9$ years (5% of the colonies in this population may reach this age, assuming the foundress queen survives that long, or colonies can requeen after death of the foundress queen).

***10-year survey of 25 colonies*:** Mintzer (2018) further reports for the survey in winter 2015/2016 that "nine colonies noted in 2006 were still extant in 2016" (page 37). Because the total number of live colonies found in the 2005/2006 survey was 25 (page 37), Mintzer's reported counts indicate that 16 of these 25 colonies died within the next 10 years, and 9 colonies were found alive in the 2015/2016 survey. Nine surviving colonies after 10 years out of 25 initial colonies indicates an estimate of an annual mortality rate of $\mu = -\ln(n_{final}/n_{initial})/(\#\text{ of years}) = -\ln(9/25)/(10) = 0.10217$. Assuming a constant mortality rate, the estimate of the average life expectancy is $1/\mu = 1/(0.10217/\text{year}) = 9.8$ years; and the estimate of maximum colony longevity is $\ln(0.05)/(-\mu) = \ln(0.05)/(-0.11123) = 29.3$ years (5% of the colonies in this population may reach this age, assuming the foundress queen survives that long, or colonies can requeen after death of the foundress queen).

The above estimates of annual colony mortality, average life expectancy, and maximum colony longevity are remarkably consistent between the 20-year survey spanning 1985 to 2005/2006 and the 10-year survey spanning 2005/2006 to 2015/2016. Specifically, the annual colony mortality during the first 20 years (low estimate of $\mu = 0.10007$; high estimate of $\mu = 0.11123$) of the 30-year survey was very similar to the annual colony mortality during the last 10 years ($\mu = 0.10217$) of the 30-year survey.

Mintzer (2018) concludes that "several *A. mexicana* colonies have survived at the national monument for more than 20 years, but none has survived over the complete 30-year survey period from 1985 to 2016" (page 37). Mintzer (2018) argues that any colony disappearance is due to mortality, not due to migration, because "colony migration (where a colony disappearance is associated with appearance of another colony 50–200 m away) is rare; I found colonies remained in the same place during visits between the decadal surveys" (page 37).

**Moser (1984).** Regarding nest longevity of *Atta texana*, Moser writes (pages 47 & 48): "According to reliable reports of rural residents, some nests may be 50 years old or more, but there is some question if the colonies occupying these nests are that old. It is doubtful if colonies live more than 11 years, because none of the queens in our laboratory colonies have exceeded that age. Although many colonies of *A. texana* have multiple queens, the new queens are accepted only when colonies are young (Moser and Lewis, 1981). Hence, there is presently no data to support the hypothesis that colonies could exist for more than 10 years by adding queens serially. Thus, the problem of how nests are able to grow old remains unresolved."

**Mueller, Ulrich (unpublished data on *A. texana* queen longevity).** A 1-year-old colony with a single queen was collected in spring 2015 in southmost Texas, then kept at Mueller's home. The colony was kept relatively small (no more than three 1-liter fungus gardens). At the time this publication went to press, the queen was alive and healthy, somewhat older than 12 years, and her colony and fungus gardens were likewise healthy.

**Mueller, Ulrich (unpublished data on longevity of *A. texana* colonies in the field in central Texas).** At the time of this publication, five large *A. texana* colonies that Mueller first saw in 1999 when he first studied the natural history of *A. texana* are still alive and well, each colony still located at the same spot where a colony were first seen in 1999. Each of these colonies was observed almost every year between 1999-2026, but these observations were not collected to quantify colony longevity (the continual presence of a live *A. texana* colony in a specific spot was noted during recreational hikes or during field work aimed at other ant species). Because each colony was already very large in 1999 (suggesting a minimal colony age of 5-10 years at that time), each of these colonies must be older than 30 years in 2026, possibly much older.

**Perfecto and Vandermeer (1993).** Perfecto and Vandermeer surveyed transects along pre-existing footpaths crossing four different habitat types (an initial survey in 1989; then a second, final survey two years later in 1991) and censused *Atta cephalotes* nests found in "an area approximately 5 m on either side of the trail" (page 381). Because both medium-sized (establishing) and older (established) colonies of *Atta cephalotes* were included in their study, Perfecto and Vandermeer (1993) write that their modeling of maximum colony lifespan predicted to be attained by 1% of the *A. cephalotes* colonies in the population yields an "extremely rough" estimate of "approximately 16 years" (page 321) maximum colony lifespan. Perfecto and Vandermeer do not calculate average colony life expectancy for their study population, but an approximation of the average colony life expectancy can be deduced from Table 1 in Perfecto and Vandermeer (1993, page 318): of the cohort of 74 *A. cephalotes* colonies included in July 1989 in the 2-year study and re-surveyed again in July 1991, 33 colonies disappeared during the 2-year observation period, and 41 colonies survived until the final survey in 1991, suggesting an estimate of annual mortality rate of $\mu = -\ln(41/74)/(2\text{ years}) = 0.2953$; and an estimate of average colony life expectancy is therefore $1/(0.2953/\text{year}) = 3.4$ years. The estimate of annual mortality rate includes both colony death and colony migration as causes of colony disappearance. The study by Perfecto and Vandermeer (1993) was not designed to estimate colony-migration rates, so it is impossible to deduce the true average colony life expectancy from the data in Table 1 of Perfecto and Vandermeer, except that the life-expectancy estimate of 3.4 years is an underestimate if some of the surveyed colonies migrated, and that the average colony life expectancy for established colonies appears to be greater than 3.4 years and smaller than the maximum lifespan of approximately 16 years estimated by Perfecto and Vandermeer.

**Roces, Flavio, personal observations of queen lifespan (until queen death) in his laboratory at the University of Würzburg**: *Atta cephalotes* (1 colony, lifespan of 13 years in lab); *Atta colombica* (3 colonies, lifespans of 7, 16, and 17 years in lab); *Atta laevigata* (7 colonies, lifespans of 7, 8, 8, 10, 10, 10, and 11 years in lab); *Atta sexdens* (2 colonies, lifespans of 8 and 15 years in lab); *Atta vollenweideri* (6 colonies, lifespans of 8, 9, 10, 11, 11, and 12 years in lab). All lab colonies were maintained healthy and well-fed; each colony had a total fungus-garden volume of at least 15 liter distributed across at least 8 garden chambers. As of April 2026, Flavio Roces also maintains live colonies of various ages in his lab: *A. laevigata* (1 colony, 10 years old); *A. sexdens* (4 colonies, each 10 years old); *A. vollenweideri* (3 colonies, each 11 years old).

**Salzmann and Jaffe (1990).** In their discussion of territoriality of *Atta laevigata* and nest-site selection of newly-mated *Atta laevigata* queens, Salzmann and Jaffe write that "… an *Atta* nest occupies the same site for several decades. A nest that recently died will soon be occupied again", presumably because newly-mated *A. laevigata* queens will found their nests at the nest site of the deceased colony (page 352).

**Weber (1972).** Weber (1972, page 38) used Kerr's (1961) estimate of the number of sperm stored in a newly-mated female of *A. sexdens rubropilosa* to calculate that the number of workers that a queen can produce during her lifetime is 2-5 million workers; that queens therefore "may have an active egg-laying life of 10-20 years"; and that, because "only one queen is known in the Brazilian mature nests of *A. sexdens*", maximum colony lifespan of *A. sexdens rubropilosa* therefore would be correspondingly 10-20 years. On page 46, Weber (1972) writes in a paragraph on *Atta cephalotes* that "one would expect the *Atta* queen to live 5-10 years."

**Weber (1976).** "A Trinidad colony of *Atta cephalotes*, originating from a May 1965 nuptial flight and removed for laboratory study on 15 July [1965], survived for 10 yr with the same queen" (page 825) in Neil Weber's lab at Swarthmore College, then the colony was moved in July 1974 to a lab at Florida State University requiring about one month of indirect transit. "Had a direct transit been possible, much damage to the delicate gardens … could have been avoided" (page 825). The colony declined around the time of the move between the two labs: "During the last colony year (July 1, 1974 – July 1, 1975) the garden volume declined slightly before it left the Swarthmore laboratory, declined further during the summer travel, … then started do decline consistently. … The number of ants declined markedly. … No brood was apparent. … The queen was clearly dead by the beginning of July" (page 828) in 1975. "The original queen survived fur 10 yr and produced no sexual brood. She produced the full range of workers" (page 827). In the Swarthmore lab, the colony was allowed to grow to an estimated volume of 11.7 liter total fungus garden in multiple garden chambers. Table 1 on page 826 of Weber (1976) shows increase and decrease of total garden volume from 1965-1975. It is unclear from Weber's account whether the stressful move between labs in July 1974 caused the eventual demise of the queen and colony within the subsequent year, or whether the queen would have died naturally anyway. The queen lived from the time of her mating flight in May 1965 until July 1975, 10 years and 2 months, that is 10.17 years.

**Weber (1982).** "Once the [*Atta* queen] dies the colony disintegrates approximately in the year her last brood lives. … Queens of *A. cephalotes* are known to have lived 10 and 12 years. It is probably that they can live 15 years in nature or in the laboratory under normal conditions and 20 years may be possible. The survival of a nest may differ from that of the original queen (plus 1 year for her last workers) if a new and young queen could develop on the periphery. Also possible is the adoption of a new queen by the colony soon after the old [queen] has died" (page 289).

**Weiss (1990).** A colony of *Atta cephalotes* was maintained by Burton Weiss in his lab at Drexel University for 114 months, that is 9.5 years. The colony was collected in 1976 (unclear which month) in Panamá as a young nest with two garden chambers. Weiss (1990) reports that the queen was assumed to have founded this nest during the "previous Spring's nuptial season" (page 422), but it is unclear from the writing whether this refers to spring 1976 or spring 1975, but most likely spring 1975, for two reasons: First, in Figure 2 on page 424, a detailed record of monthly foraging by the colony starts in July 1976 and ends in October 1985 (suggesting a collection date in the first half of 1976, and hence the colony was apparently founded in spring 1975); and second the colony was collected with two gardens in 1976 (too large if the colony had been founded in May 1976); these combined facts suggests that the colony must have been founded in spring 1975, giving it sufficient time to grow two gardens by the time of collection in early 1976. Weiss carefully documented colony growth and eventual decline of the colony: "the queen died in mid-October 1985, having produced no brood since May 1985, and the last workers expired on December 18, 1985" (page 428). Figure 2 on page 424 of Weiss (1990) shows that the decline of the colony was manifested also in the gradual decline of foraging activity starting in May 1985. Assuming that the queen founded the nest in May 1975 (May is the typical month of mating flights for *A. cephalotes* in Panamá, hence the name "zompopos de Mayo" for *A. cephalotes* alates), the queen lived for 10 years and 5 months from May 1975 until October 1985 (10.4 years). Although the queen died in mid-October 1985, she sustained the colony with brood only from May 1975 until May 1985, a time period of exactly 10 years (longevity as a functional queen). Unlike most *Atta* colonies maintained in other labs, the colony was allowed to grow numerous gardens for a total volume of up to 50-55 liters between all gardens. The colony produced soldiers, but never produced alate reproductives. The queen produced many more workers than in typical lab colonies, but far less than in an established colony in the wild. It is possible that the queen ran out of sperm in May 1985 and therefore was no longer able to sustain worker production, but other factors such as disease can also explain why the queen stopped producing workers in 1985.

**Wirth et al. (2003).** Table 9 on page 87 in Wirth et al. (2003) reports, for a cohort of 51 established *A. colombica* colonies observed from May 1996 until October 1998 (17 months total) on Barro Colorado Island, Panamå, that 13 of these colonies had died by October 1998. Wirth et al. calculate from their data in Table 9 a mortality rate of 0.092/year (Table 9 in Wirth et al. 2003). Assuming constant mortality, and taking the mortality estimate of $\mu = 0.092$/year reported in Table 9 by Wirth et al., this mortality rate corresponds to an average colony life expectancy for established *A. colombica* colonies of 1/(0.092/year) = 10.9 years. The exact survey methods of Wirth et al. were: "A detailed survey was performed in 1996. Subsequently, colony dynamics were recorded from May 1996 until October 1998 in 3-month intervals during the first year, and monthly during the rest of the investigation period. Nests were regarded as newly established when they had grown to sufficient size to be included in the survey. … Colonies were regarded as dead when there were no visible ant activities at the nest site for several weeks and also no signs of colony movement" (pages 84 & 85).

Cole (2009) reanalyzed the demographic data reported by Wirth et al. (2003), but reports the methods of Wirth et al. as "survival from multiple censuses, N = 92 colonies, 2 years". We could not reconstruct these exact data from Table 9 in Wirth et al., but we found on page 85 of Wirth et al. that the total number of different *A. colombica* nests known to Wirth et al. across the entire island was 92 colonies (these included colonies that were alive from the beginning until the end of the survey period; colonies alive at the beginning of the survey period but dying sometime during the survey period; and newly-founded colonies not known at the beginning of the survey period and presumed therefore to be newly founded sometime during the survey period), and we found in the same paragraph on page 85 also a reference to a survey period "between 1996 and 1998", a period of 2 years. It is possible that Cole (2009) used these data of 92 colonies and a survey period of 2 years total in his calculations, whereas the survey period covered in Table 9 of Wirth et al. was from May 1996 until October 1998, covering only 17 months total, not 2 full years (i.e., Cole apparently did not reanalyze the data in Table 9, but a different, somewhat larger dataset that we are unable to reconstruct from the publications). Because we were unable to reconstruct the exact calculations of annual mortality rates reported by both Wirth et al. and by Cole, we list in our Table S1 the mortality rates exactly as reported by Wirth et al. (2003) and Cole (2009).

Interestingly, per-colony migration rate was estimated by Wirth et al. to be 25% per year for the same population (Table 9 in Wirth et al. 2003). If nest disappearances due to migration had been incorrectly judged to be cases of colony mortality, an annualized mortality rate of $\mu$ = 0.092/year + 0.250/year = 0.342/year would have been incorrectly calculated for this population, leading to an under-estimate of average colony life expectancy of only 1/(0.342/year) = 2.9 years. This illustrates the importance of accounting for colony migrations when estimating life expectancies of *Atta* colonies. It is possible that Perfecto and Vandermeer (1993; see above) estimated for established colonies of *Atta cephalotes* at La Selva, Costa Rica, a likewise high annual mortality rate of $\mu$ = 0.2953/year [corresponding to an average colony life expectancy of only 1/(0.2953/year) = 3.4 years], because Perfecto and Vandermeer did not account for colony migration as one possible cause of colony disappearance.

**THE FOLLWING ARTICLE IS NOT INCLUDED IN TABLE S1, BECAUSE MANY CLAIMS IN THIS ACCOUNT ARE SPECULATIVE, UNRELIABLE, OR OUTRIGHT FALSE:**

**Schade F. H. (1973).** Schade writes on page 86: “Paraguayan farmers usually say 30 to 40 years [of lifespan of an *Atta* nest]. … A 92-year-old man showed me a nest beside a house where he was born. The nest was just as active as it had been in his childhood. A conservative estimate of the age of this colony would be 130-140 years. Probably it is much older. Certainly all nests do not reach such an advanced age, but at least it is possible.” Based on this communication from the 92-year-old man, Schade argues that the lifespan of an *Atta* nest may sometimes exceed 100 years. Schade further considers the possibility of satellite mounds (Schade calls these “branch offices”) associated with a main mound, and that overall longevity of such a polydomous colony maybe be older than estimated from a mix of longevity estimates of central mounds and the more ephemeral satellite mounds. Schade also cautions that colony migrations may complicate longevity estimates of nest-site-switching colonies. Because Schade believes that a single queen lives maximally for about 15 years, Schade speculates that the much longer longevity of colonies can be explained only if one assumes that colonies re-queen, and, because of the possibility of queen adoption during the lifetime of a nest, there may be multiple queens per *Atta* nest. Fowler et al. (1986) write disparagingly of Schade’s (1973) suggestion that maximum lifespan of *Atta sexdens rubropilosa* may exceed 100 years, adding this biting comment: “Schade (1973) referred to colony life spans of *Atta sexdens rubropilosa* on the order of 30 to 140 years, but apparently these estimates are not based on data” (Fowler et al. 1986, page 128). We perused Schade (1973) carefully, and we agree with this assessment by Fowler et al. (1986).

**ALSO NOT INCLUDED IN TABLE S1:**

**Nogueira et al. (2022).** Nogueira et al (2022) write that “*Atta* queens are known to live for more than 20 years” and Nogueira et al cite here Autuori (1941; Arquivos do Instituto Biologico São Paulo 12: 197-228), but we were unable to find a statement of 20-year queen longevity in Autuori (1941).

### DETAILED METHODS

The following is a detailed account of our Methods; these are the same Methods that we summarize concisely in the main article:

**Field Surveys:** Our estimates of *A. texana* colony longevity are based on two surveys: (i) a 19-year survey of 9 established *A. texana* nests that we first geo-referenced in Louisiana in 2006 for population-genetic studies of the ant-cultivated fungi (Mueller et al. 2011a,b) and that we re-surveyed again in 2019, 2020, and annually in 2022-2025; and (ii) a 2-year survey of 195 established nests surveyed annually in 2023, 2024, and 2025.

**Size and appearance of *A.texana* nests at the start of surveys:** Each *A. texana* nest included in our surveys had (i) a mound diameter of at least 7 meter, and (ii) extensive recent accumulations of fresh excavate on the central mound (Figure 1) suggesting healthy worker populations. Following the methods of Cahal (1993), we recorded at each visit the dimensions of the contiguous area of excavated soil and excavation crater defining the central mound, estimating in meters the length of the mound along its longest axis, and the breadth of the mound perpendicular to that first axis. We timed the surveys to occur during periods with sufficiently warm days when workers excavate soil, and to occur at least one week after rains, such that the recent accumulation of fresh excavate at excavation craters on the mound indicated that a colony was alive and well. Presence of excavating or foraging workers on the mound is a direct indication that a colony is alive, but workers become inactive on the mound on hot days or cold days, and at such times we used the presence of fresh, pelleted excavate (i..e, soil pellets what were brought by workers to the surface and discarded on the mound very recently) as a proxy to evaluate whether a specific *A. texana* colony was alive.

**Definition of "established nest":** We refer to each nest included in our survey as an "established nest", that is, an *A. texana* nest that had reached a mound size of at least 7 meter diameter and therefore had survived the high-mortality phase of smaller, young *Atta* nests typical for Type III survivorship (Autuori 1950; Mariconi 1970; Fowler et al. 1984, 1986), and typical for ants at large (Cole 2009; Vergara-Martínez et al. 2024).

Because we biased our samples to include only large established *A. texana* nests (i.e., we excluded nests of less than 7 meter mound diameter), our estimates of nest mortality and nest longevity are not demographic estimates for average *A. texana* nests in a population, but estimates for a subset of *A. texana* nests, specifically those nests that (i) had survived the high-mortality phase of smaller and younger nests; (ii) had therefore become established; (iii) had larger foraging territories and were therefore more likely found by us; and (iv) had conspicuous mounds.

**The age of each *A.texana* colony at the start of our surveys was likely at least 5-10 years, possibly much older:** The exact ages of *A. texana* colonies at the start of our surveys were not known. Because we included in our surveys only large colonies with above-ground mounds of at least 7 meter diameter, we estimate that each colony must have been at least 5-10 years old at the start of each survey. It is possible that the average colony included in our study was actually much older, because the nests included in our surveys measured typically 10-15 meter mound diameter (range 7-35 meter mound diameter). Our minimal estimate of at least 5-10 years colony age at the start of our surveys is supported by the following arguments and observations:

(a) Mound diameter is a proxy of colony age, not an exact measure of age; it does not correlate linearly with colony age in *A. texana*. As in other *Atta* species (Autuori 1941, 1942; Mariconi 1970; Hernández et al. 1990), and based on our own observations, mound diameter and colony age in *A. texana* correlate more closely at younger ages (we estimate when a colony is less than 5-10 years old) and less closely at older ages (more than 15-25 years).

(b) In *A. laevigata* (a species that has similar nest architecture as *A. texana*, constructing similar extensive tunnel systems around its mounds in Venezuela), mound area correlates closely with colony age during the first 4 years, but "the rate of increase of the nest area was higher during the first 30 months than during the final two years. Colonies with 4 years of age had nests with an average size of 71 ± 13 $m^2$" (Hernández et al. 1990), which corresponds to approximate mound dimensions of 8.4 meter X 8.4 meter.

Because colonies of tropical *Atta* species grow faster in size under the warmer underground and above-ground temperatures of tropical habitat than colonies of *A. texana* in their cooler subtropical and temperate habitats, and because 4-year-old nests of *A. laevigata* grow to an average size of 8.4 meter X 8.4 meter in tropical Venezuela (Hernández et al. 1990), a mound size of 7 meter X 7 meter diameter in *A. texana* suggests an age older than 5 years old for such colonies. This supports our approximation that *A. texana* mounds of 7 meter X 7 meter diameter are likely at least 5-10 years old.

(c) Unlike for *A. laevigata*, the correlation between mound diameter and colony age has not been quantified for *A. texana*. Our understanding of the relationship between mound diameter and colony age is based on *A. texana* colonies that we have observed continuously in central Texas during 27 years of field research. Colonies that migrate (relocate between different nest sites; see below) can undergo a reduction in mound size at the time of migration (i.e., the new mound after migration tends to be somewhat smaller than the mound before migration). Because some *A. texana* colonies migrate several times during their lifespans within their territories, migration is one reason why colony age and mound size is less correlated at older ages in *A. texana*. Fortunately, only about 2-3% of established colonies migrate every year in Louisiana (see below).

(d) During field work, it was easiest to find large mounds because of their corresponding large foraging territories. Specifically, we typically found first one of a colony's distant foraging exits (so-called feeder exits, linked to a colony's central mound by long underground tunnels up to 150 meter long, but more typically 20-80 meter long), before we found the colony's central mound. Moreover, we located about 20% of the *A. texana* colonies included in our study because they had exceptionally large and conspicuous mounds that were noticeable on the satellite images of Google Maps. Our sample of surveyed nests is therefore very biased towards colonies with large foraging territories and large conspicuous mounds, and correspondingly likely biased also towards very old colonies (i.e., young *A. texana* colonies were less likely included in our surveys because of their smaller, less conspicuous mounds and their smaller foraging territories).

**19-year survey of 9 *A.texana* nests:** Our first sample of 9 nests of the 19-year survey included all the *A. texana* nests that we had geo-referenced in 2006 in Louisiana. Four of these 9 nests we dug into in 2006 to reach the topmost fungus gardens (1.4–3.1 meter deep) to obtain live isolates of the ant-cultivated fungi for quantitative-genetic experiments and population-genetic analyses (Mueller et al. 2011a,b); these four nests are indicated in Table 2 and Table S1 with asterisks. Because of the large sizes of *A. texana* nests (fungal gardens occur 1.5–8 meter below ground in a nest; queens are thought to inhabit the deeper fungal chambers), digging a hole that affects less than 10% of the mound and that reaches only the topmost fungal gardens, then refilling the hole after garden collection, should not have affected colony survivorship, but a minor impact on survivorship cannot be ruled out completely.

**2-year survey of 195 *A.texana* nests:** Our second sample of 195 established nests of the 2-year survey were selected from 301 nests geo-referenced during annual field work from 2019 to 2023 in Louisiana. We excluded from these 301 nests 106 nests because they were in locations with high likelihood of disturbance (e.g., yards of houses, old pine plantations approaching the time of logging), or nests where we would not be able to search the surround within a 100-meter radius in all directions (e.g., we excluded nests located next to a fenced-in, private property). Six of the 9 nests included in our 19-year survey were also included in our 2-year survey (2 nests in the 19-year survey had died by the time the 2-year survey was started in 2023, see Table 2 and Table S1; and one nest in the 19-year survey was at the edge of fenced-in private property, and the area surrounding this nest therefore could not have been searched within a 100-meter radius in all directions to differentiate between death and migration of that colony if the mound was found unoccupied by ants at the time of a re-survey; see below).

**Access to *A. texana* nests included in the 2-year survey:** Accessibility of a 100-meter radius around a nest was important for re-surveys to distinguish between migration versus mortality as causes of any colony disappearance if a mound was found unoccupied by live ants at the time of a re-survey. Both a dead mound and a mound abandoned by a migrating colony can look very similar (absence of workers, no fresh excavate, and no open exits on a mound), but an *A. texana* colony that migrates has active foraging exits with fresh excavate at regular intervals between the abandoned mound and the new mound (sometimes a few active exits remain for years on the abandoned mound, and the abandoned and the new mound are linked by underground tunnels), whereas a colony that dies has no active foraging exits nearby its uninhabited former mound. Fortunately, colony migrations in Louisiana are rare, and the rate of colony disappearance is low (see Table 2 in main article), so we had to conduct in re-surveys few careful searches within a radius of 100 meter of any inactive mound to distinguish between migration and death, using the above criteria. For most nests we simply had to revisit a nest's GPS location of a previous survey to verify that a colony was still alive at that same GPS location, and we could infer from the size and the growth trajectory of the mound that the colony was well (i.e., the mound size stayed the same or increased compared to the mound size recorded in the previous survey; there was no drastic reduction in mound size, as occurs in moribund nests, see below).

Finding *A. texana* nests with a 100-meter-radius access and with potentially high human impact (i.e., nests in unstable habitat; e.g., in cemeteries) was more difficult than finding nests with 100-meter-radius access and little human impact (i.e., nests in more stable habitat; e.g., in pine plantations). Consequently, our sample size of nests in the Control Sample (n=24 nests in unstable habitat, where we expected human impact to be more frequent) is much smaller than the sample size of nests in the Main Sample (n=171 nests in stable habitat, where we expected human impact to be less frequent; see below).

**Baseline survey in January 2023, re-surveys in January 2024 and January 2025:** The 2-year survey consisted of a baseline survey in January 2023, then re-surveys of all nests (including nests presumed dead or moribund) in January 2024 and January 2025. About 37% of the nests in the January 2023 baseline survey were actually surveyed one month earlier in December 2022, and for these nests the between-survey timespan from initial survey in December 2022 to re-survey in January 2024 is 13 months; but to simplify analyses, we pool these nests with the nests surveyed in January 2023, to calculate mortality rates on an annual basis rather than per-month basis. The 2-year survey therefore covers a time window of 25 months for 73 nests (37% of 195 nests total) surveyed from December 2022 to January 2025, compared to a time window of 24 months (i.e., exactly 2 years) for 122 nests (63% of 195 nests total) surveyed from January 2023 to January 2025. The 73 nests surveyed across a 25-month time window were all located in the southern half of the *A. texana* range in Louisiana (Table S3), and included 10 nests in the Control Sample (41.7% of the 24 nests total in the Control Sample), and 63 nests in the Main Sample (36.8% of the 171 nests total in the Main Sample). Pooling the nests surveyed for 25 months with the nests surveyed for 24 months should not bias our estimates of colony mortality, because (i) mortality rates were overall very low; (ii) mortality rates in the southern half of the *A. texana* range in Louisiana did not differ from the mortality rates in the northern half of Louisiana (i.e., nests in both the southern and northern half of Louisiana died or became moribund in about equal proportions); and (iii) nests in the Control Sample (higher mortality rate than nests in the Main Sample; Table 3) were distributed approximately equal between the southern and northern half of the *A. texana* range in Louisiana.

Many of the 195 nests in the 2-year survey had been found during field work in years before the baseline survey in January 2023, seven of these nests dating back to our very first field work in Louisiana in 2006, but most nests had been found between 2019 and January 2023 (Table S3). By January 2023 (and including new nests found in the survey in January 2023), we had geo-referenced a total of 301 established nests of *A. texana* in Louisiana. In December 2023, before the first re-survey in January 2024, we selected from these 301 nests for our 2-year survey 195 established nests with accessibility of a radius of 100 meter around each nest (see above). Specifically, we excluded from the 301 nests 106 nests because they were in locations with high likelihood of disturbance (e.g., yards of houses, old pine plantations that we judged were approaching the time of logging), or nests where we would not be able to search the surround within a 100-meter radius in all directions (e.g., nests located next to a fenced-in property). Accessibility of a 100-meter radius around a nest was important for re-surveys to distinguish between migration versus mortality as causes of any colony disappearance (see above). All field records and measurements of the 195 surveyed mounds, including records from surveys preceding the years before 2023, are in Table S3.

**Assignment of nests to Control and Main Samples:** Of the 195 established nests in searchable habitat included in our 2-year survey, we assigned, before the first re-survey in January 2024, 171 colonies to a Main Sample because these nests were judged to be in more stable and more protected habitat where we thought disturbance by humans would be infrequent (e.g., National Forest; younger pine plantations;

semi-protected areas along gas pipelines or under powerlines). For comparison, we assigned the remaining 24 nests to a Control Sample because we judged these nests to be in less stable and less protected habitat (e.g., in cemeteries, right at or under paved roads) where we expected disturbance by humans to be more frequent (e.g., by poisoning or physical disturbance of *A. texana* nests). We assigned the 195 nests to Control Sample versus Main Sample in December 2023 by considering information recorded in our field notes, and by inspecting the nest locations at Google Maps.

Because assignments to Control versus Main Samples were made before our first re-survey in January 2024, the assignments were pre-planned, and therefore independent of later observations of mortality or morbidity of surveyed nests.

**Identification of live, moribund, and dead *A. texana* nests:** Each time a nest was re-surveyed, we recorded the visible signs that a nest was inhabited by a live *A. texana* colony, such as (i) excavating workers observable on the mound surface, and, if workers were not active on the mound during very hot or cold times, the level of recently accumulated fresh excavate on the mound (ranging from no fresh excavate to very much fresh excavate); (ii) the extent of well-sculpted thatching built by the ants to cover exits on a mound; (iii) clipping by the workers of encroaching vegetation to keep the mound free of plant growth; and (iv) presence of active (currently-used) foraging exits in the colony's foraging territory surrounding its mound. Following the methods of Cahal (1993), we also recorded at each visit the dimensions of the contiguous area of excavated soil and excavation crater defining the central mound, estimating the length of the mound along its longest axis, and the breadth of the mound perpendicular to that first axis. This combined information allowed us to classify each nest into (a) *alive* (presence of workers, fresh excavate, well-thatched exits, or recently cut vegetation on the mound); (b) *dead* (no workers, no fresh excavate, no open exits, no thatched exits, only washed-out eroded excavate on the mound, no active foraging exits nearby); or (c) *moribund* (noticeable reduction in mound dimensions between preceding survey and re-survey; little or minimal fresh excavate on mound at the time of the re-survey; some worker activity evident indicating no complete absence of live *A. texana*, such as actively excavating workers found on the mound, little or minimal fresh excavate on the mound, or well-maintained, open foraging exits within a 30 meter radius of the mound).

To verify the status of the dead nests (n=7) and moribund nests (n=2) observed in our second re-survey in January 2025 (Table 2), we revisited these nine nests in mid-April 2025. April is the time in spring when mature mounds prepare the mound for mating flights in late April or May, and April is therefore one ideal time to evaluate the status of an *A. texana* colony. Signs of colony health and vigor in spring include (i) workers cut most of the vegetation growing on and at the periphery of the mound, to enable reproductive alates to depart readily for their mating flights; (ii) the cut vegetation accumulates as fresh plant material (e.g., leaf discs) on the mound; and (iii) nest entrances on the mound are thatched by the workers with this freshly cut plant material, such that workers can remove the thatch before a mating flight, then re-thatch exits quickly after completion of the mating flight, thereby opening and closing the nest exits). April is therefore an ideal time to assess whether an *A. texana* colony is alive and healthy, moribund, or dead. All nests judged in January 2025 to be dead (n=7) were confirmed dead in April 2025; and all nests judged in January 2025 to be moribund (n=2) were confirmed moribund in April 2025 (see details summarized for each of these nests below).

**Modeling of life expectancy and maximum lifespan:** We used the same approach of estimating annual mortality, life expectancy, and maximum life expectancy that we used to model these parameters in the primitive fungus-growing *ant Mycetosoritis hartmanni* (Mueller et al. 2023). As in other studies of ant colony longevity (e.g., Pamilo 1991; Meyer et al. 2009; Cole 2009), this approach assumes that mortality is independent of age for established colonies (i.e., colonies that had survived the high-mortality stage of young colonies), and longevity can be estimated under a constant hazard model (i.e., colonies die off at a constant rate after establishment). From the observed survivorship records in our 19-year survey (n=9 nests) and 2-year survey (n=171 main nests, n=24 control nests), we derived estimates of the annualized mortality rate (μ, mortality per year) of nests. In both our 19-year and our 2-year multi-year survey, the number of surviving colonies declines every year due to mortality, so we estimate mortality across multiple years as $\mu = -\ln(n_{final}/n_{initial})/(\#\text{ of years})$ [solving for μ from: $n_{final} = n_{initial} * e^{-\mu * time}$], where $n_{final}$ is the number of nests alive at the end of a multi-year survey, $n_{initial}$ is the number of nests alive at the beginning of the multi-year survey, and *# of years* is the total number of years elapsed between initial and final census. Assuming that mortality is constant with age for established colonies (see above), we use our estimate of mortality (μ) to calculate (i) an estimate of the expected lifespan (1/μ, the inverse of μ; i.e., the mathematical expectation of an exponential distribution); (ii) an estimate of the expected "maximum" lifespan, defined as the age of the nests when less than 5% of an initial cohort is expected to have survived [$-\mu^{-1} * \ln(0.05)$, in years]; and (iii) the range of this expected maximum lifespan derived from our estimate of mortality ± standard error of a binomial distribution [$1/(\mu+SE_{\mu})$ to $1/(\mu-SE_{\mu})$, in years; where $SE_{\mu} = [\mu * (1-\mu)]^{0.5}/n_{initial}$) and $n_{initial}$ is the total number of nests at the beginning of a survey]. Constant annual colony survival likely approximates the true survival rate for much of the life of established colonies. In *A. texana*, colony survivorship does not appear to decline markedly in established nests after the death of the foundress queen(s) because large established nests appear to adopt new queens regularly (see Results and Discussion).

### COLONY MIGRATION

**Different mature colonies of *A. texana* in Louisiana are spaced at least 110-150 meter apart (Figure 1):** Overdispersion of mature *A. texana* colonies, as in Figure 1, is one reason why it is possible to differentiate between migration and colony death as reasons for colony disappearance. In Louisiana, different neighboring colonies maintain a minimal distance of about 110-150 meter between nest mounds (Ulrich Mueller, unpublished data) because of territorial interactions. Each colony builds underground tunnels (typically 10-80 meter long, but up to 150 meter long in Louisiana) that radiate out from its central mound, and above-ground foraging trails starting at tunnel exits (so-called foraging or feeder exits) radiate out even further. The contact zone between neighboring colonies is a zone where foraging exits are absent (i.e., the tunnel systems of different colonies do not connect). That is, when surveying the ground in a straight line between two neighboring *A. texana* mounds, the density of foraging exits initially decreases when walking from one mound to a neighboring mound, then there is a zone where foraging exits are absent, then the density of foraging exits gradually increases when approaching the neighboring mound. The presence of an intermediate zone free of foraging exits can be used as an indicator that two neighboring *A. texana* mounds are not linked by underground tunnels.

Sometimes two large *A. texana* mounds can be found relatively close to each other, and in such cases there are foraging exits at regular intervals and at high densities between these two mounds (i.e., there is no zone between these two mounds where foraging exits are absent), indicating that these two mounds are linked by underground tunnels. Cases where two mounds are found in close proximity are typically cases where an *A. texana* colony is in the process of migrating from one mound to the other mound, and during this transitional period a colony is temporarily polydomous (a colony has two interconnected nest sites) as the ants are relocating brood and fungus gardens in their underground tunnels from pre-migration nest site to post-migration nest site. Unlike in tropical *Atta* species, colony migrations of *A. texana* colonies have never been observed above ground; instead, *A. texana* uses its underground tunnels when relocating between nest sites.

If a colony has completed its migration between nest sites, the colony typically maintains active exits at the pre-migration nest site for some time (sometimes active exits are maintained for years at the pre-migration nest site) and active foraging exits persist between pre-migration and post-migration nest sites (i.e., the colony maintains the underground tunnels used for migration between old and new nest). In contrast, if a colony dies, there are no active, well-maintained foraging exits nearby the dead nest, and any signs of former foraging exits and foraging crater disappear within a few months as the foraging crater become eroded by rain and wind, become covered by leaves, or become overgrown with vegetation. A moribund colony, a stage between alive and dead colony, can be identified because a moribund colony shows (i) a marked reduction in mound size between surveys; (ii) some active foraging exits in the near surround of the mound (e.g., ≈25 meter surround), but fewer and fewer exits at greater distances from a mound; (iii) few active open exits on the mound, and (iv) most former exits on the mound are eroded and closed. A moribund nest can be distinguished from an abandoned pre-migration mound after colony migration in that a moribund nest has no active foraging exits beyond ≈25 meter distance from the mound (these foraging exits can be active for months in the case of moribund mounds, until all the workers in the moribund colony have died), but in the case of an abandoned pre-migration mound, there are continuous foraging exits between pre-migration and post-migration mounds, and these foraging exits can persist for years between pre-migration and post-migration mounds as the colony maintains underground tunnels between pre-migration and post-migration nest sites. In many cases of migration (see below some such observations from our study), a colony that migrates in one year to a new site may migrate back within the next year to its original pre-migration nest site. We know of some *A. texana* colonies that we have observed in central Texas for a decade and that alternate (toggle) back-and-forth almost annually between two nest sites. It appears that three of the *A. texana* colonies included in our Louisiana study may also be such toggling colonies (details below).

**Migrations of *A. texana colonies* are very rare in Louisiana:** In addition to the aforementioned overdispersion of *A. texana* nests in Louisiana, the rarity of colony migrations by large established colonies in undisturbed habitat in Louisiana is a second reasons why we decided to conduct this colony-longevity study in Louisiana. Because the center of an *Atta* mound can shift slightly between years whenever nests add new underground fungal chambers primarily at one end of the mound, and because of the accuracy of our GPS measurements was typically 3-5 meters (Table S3), we regard any difference in GPS location of a mound between survey years of less than 20 meter not as a true colony migration, but as a natural minor shift typical for all *A. texana* nests. Of the 195 nests in the 2-year survey, twelve nests migrated more than 20 meters within the 2-year survey window [11 nests (6.4%) in the Main Sample of 171 nests; and 1 nest (4.2%) in the Control Sample of 24 nests]. Most nests that migrated did so only once during the 2 years, but 3 colonies migrated to a new location in the first year, then migrated back to the pre-migration site in the second year (so-called toggle colonies). Such migrations back-and forth between different locations within the territory of a colony appear to be the most frequent kind of migrations of established *A. texana* colonies in undisturbed habitat.

Nest relocations of *A. texana* colonies are typically triggered by disturbance, such as poisoning by humans; or by physical disturbance of the nest site, for example by logging of trees surrounding a mound, or brush-clearing along roadsides or gaslines when large machinery or tractors are driven over an *A. texana* mound. Controlled-burn fires of managed pine forests sometimes can trigger colony migrations (e.g., in controlled-burn sectors of National Forest), but logging of trees close to nests appears to be a more pronounced physical disturbance, triggering colony migrations more predictably in *A. texana*. Even though we selected 171 nests in habitat were we thought disturbance by humans was unlikely, some of these nests were unfortunately disturbed by humans during our 2-year survey, for example by logging (in locations where we misjudged the age of a pine plantation, and where we had wrongly judged that logging would not occur during the two years of the survey).

The great majority of *A. texana* colonies (93.8% of 195 colonies total) remained at their baseline GPS location during the 2-year survey (i.e., 183 of the surveyed *A. texana* colonies did not migrate between survey years). The average change in GPS location for these colonies was 6.5 meter between survey years. Such small “shifts” in location can be attributed to a combination of (i) GPS-measurement error (typically 3-5 meter accuracy of our GPS device) and (ii) gradual and natural shifts of a mound center as underground chambers and corresponding excavation crater on the surface are added predominantly at one end of a mound.

Twelve colonies (6.2% of 195 colonies total) migrated or shifted their nest location for more than 20 meter between some survey years. As described below in detail, most of these migrations for more than 20 meter occurred after physical disturbance of mounds (e.g., logging of trees right at the mound, heavy vehicles were driven over a mound) or after alteration of nearby habitat (opening of habitat by controlled burns or by logging at a short distance from a mound). Our field notes of these 12 colony relocations are recorded in Table S3 and are summarized here in narrative format:

**Migrations of Main-Sample Colonies between January 2023 – January 2025 during 2-year survey:**
**Colony #0011** relocated between December 2022 and January 2024 from a location at the forest edge at the west side of a paved road to a new location in a recently logged and replanted pine plantation at the east side of that paved road, a nest-relocation distance of 50.8 meter. Colony #0011 was first found in November 2020 31.2 meter west of the location of Colony #0011 in December 2022 (i.e., Colony #0011 migrated 31.2 meter before the start of the 2-year survey to the location in December 2022 at the start of the 2-year survey). The mound was of intermediate size in November 2020 (9 meter X 8 meter), and the mound increased in size gradually until the last observation in January 2025 (16 meter X 9 meter). Already in December 2022, the beginnings of a possible satellite mound was noted ≈50 meter to the east

of the pre-migration mound, as well as active foraging exits at regular intervals between the pre-migration mound and this smaller mound (recorded as possible "satellite mound" in the field notes), suggesting that the colony migration appears to have started in late 2022. In January 2024, there were a few active exits on the pre-migration mound (i.e., the pre-migration mound was not completely abandoned), and there were active foraging exits at regular intervals between the pre-migration and post-migration mounds; this indicates that the pre-migration and post-migration mounds were linked by underground tunnels inhabited by the same *A. texana* colony. Between January 2024 and January 2025, Colony #0011 remained in its post-migration mound (shifting in the second year only 2.2 meter in GPS location). The mound sizes of Colony #0011 estimated in years after migration were larger than the mound sizes estimated before migration; this increase in estimated mound size may be an artifact of small habitat differences before and after migration, or a consequence of colony growth (increase in worker number), or a combination of both. The migration of Colony #0011 may have been triggered by the logging in 2022 of the forest to the east of the paved road, the opening of un-forested habitat there, and the planting of pine seedlings that *A. texana* prefers for leaf-cutting of pine needles to support growth of fungal gardens.

**Colony #0012** relocated between January 2024 and January 2025 from a site at the forest edge at the west of an unpaved forest road into a nearby opening in the forest to the east of that forest road, a nest-relocation distance of 61.9 meter. Colony #0012 was first found at its pre-migration site west of the unpaved road in November 2020. The mound of Colony #0012 was estimated to be relatively large (13 meter X 10 meter) in November 2020. In January 2025, the size of the mound at the pre-migration site seemed reduced compared to earlier years, and there seemed to be a reduced number of active exits on the pre-migration mound; this prompted a search for a new location nearby to which Colony #0012 may have migrated. When following a series of foraging exits to the east of the pre-migration mound (few foraging exits were found to the north, west, or south of the pre-migration mound), the post-migration mound was found 61.9 meter to the east in a small clearing in pine forest. To follow up on this migration, Colony #0012 was re-surveyed in April 2025. Between January 2025 and April 2025, Colony #0012 relocated back to its pre-migration location from January 2023 and January 2024 (i.e., the pre-migration mound was re-occupied by April 2025, and the pre-migration mound appeared like a regular mound; whereas the new, temporary mound to the east seemed to be used less by the colony), a back-migration distance of 58.1 meter. The reasons for the back-and-forth migrations are unclear, as the pine forest and the forest edges near Colony #0012 did not appear to have been disturbed during that time window.

**Colony #0017** relocated in late 2024 or early January 2025 from a site at the forest edge at the north side of an unpaved forest road to the forest edge at the south side of that forest road, a nest-relocation distance of 32.3 meter. Colony #0017 was first found at its pre-migration site in December 2022. The mound of Colony #0017 was estimated to be relatively large (14 meter X 8 meter) in December 2022, and the mound size remained approximately the same until January 2025 (15 meter X 11 meter) and April 2025 (12 meter X 11 meter). In January 2025, the site of the pre-migration mound had been greatly disturbed by recent logging (logging vehicles had been driven over the pre-migration mound; new pines had not been replanted by January 2025), and the post-migration mound (15 meter X 11 meter) was clearly visible 32.2 meter distant and across the forest road from the location of the pre-migration mound. By April 2025, Colony #0017 remained at its post-migration nest site south of the forest road, and the colony seemed healthy at its post-migration site and little affected by the logging across the forest road. The migration was undoubtedly triggered by the logging of the pine forest north of the forest road in late 2024, and Colony #0017 responded to that disturbance by migrating a short distance to the undisturbed forest edge at the south side of that same forest road.

**Colony #0044** relocated between January 2024 and January 2025 from a site at the side of a hiking trail in semi-open controlled-burn pine forest to a location 58.0 meter distant to the west, at the side of the same hiking trail. Colony #0044 was first found at its pre-migration site in December 2022. The mound of Colony #0044 was estimated to be very large (18 meter X 14 meter) in December 2022, and the the post-migration mound increased in size by January 2025 (20 meter X 19 meter). The post-migration site at the hiking trail had been surveyed during the two years preceding the migration, while searching for signs of *A. texana* along the hiking trail in December 2022 and January 2024; at both times no mound had been noticed at the future post-migration site. In January 2025, there were a few active exits with some fresh excavate on the pre-migration mound (i.e., the pre-migration mound was not completely abandoned), and there were many active foraging exits at regular intervals between the pre-migration and post-migration mounds; this indicates that the pre-migration and post-migration mounds were linked by underground tunnels inhabited by the same *A. texana* colony. The migration of Colony #0044 may have been triggered by a controlled burn of the forest in 2024.

**Colony #0048** relocated between December 2022 and January 2024 from a location at a hunter campground in semi-open pine forest to a somewhat less-disturbed location along a nearby narrow forest road (the access road to the hunter campground), a nest-relocation distance of 54.7 meter. Colony #0048 was first found at its pre-migration site in December 2022. In January 2024, there were a few active exits on the pre-migration mound at the hunter campground (i.e., the pre-migration mound was not completely abandoned), and there were active foraging exits at regular intervals between the pre-migration and post-migration mounds; this indicates that the migration and post-migration mounds were linked by underground tunnels inhabited by the same *A. texana* colony. Colony #0048 then relocated again between January 2024 and January 2025 along that same narrow forest road, a second nest-relocation distance of 27.6 meter. Colony #0048 had first been found at its pre-migration site at the hunter campground in December 2022; the mound of Colony #0044 was estimated to be very large (16 meter X 15 meter) at that time in December 2022, decreased after the first migration to 7 meter X 6 meter in January 2024, then increased after the second migration to 12 meter X 11 meter in January 2025. The first migration in 2023 may have been triggered by the disturbance by trucks or all-terrain vehicles driving over the mound during the hunting season at the end of 2022. The second migration in 2024 may have likewise been triggered by the disturbance by trucks or all-terrain vehicles driving over part of the mound in 2024.

**Colony #0056** relocated between December 2022 and January 2024 from a location at the edge of a recently logged area at the east side of a paved forest road to a new location at the edge of undisturbed forest to the north-west and across the paved forest road, a nest-relocation distance of 41.9 meter. Colony #0056 was first found at its pre-migration site in December 2022. In January 2024, there were a few active exits on the pre-migration mound and some well-thatched exits (i.e., the pre-migration mound was not completely abandoned), and there were active foraging exits at regular intervals along both roadsides between the pre-migration and post-migration mounds; this indicates that the pre-migration and post-migration mounds were linked by underground tunnels under the road that were inhabited by the same *A. texana* colony. Colony #0056 then relocated between January 2024 and January 2025 from the post-migration location back to very near the pre-migration location from December 2022, a second nest-relocation distance of 63.0 meter. The mound of Colony #0044 was estimated to be very large (15 meter X 14 meter) in December 2022 before the first migration; the mound increased slightly in size after the first migration (17 meter X 14 meter in January 2024); then the mound size returned in January 2025 after the back-migration to the initial mound size (15 meter X 14 meter). The back-and-forth migration may have been triggered initially by logging east of the forest road and in the immediate surround of Colony #0056 (and possibly heavy logging machinery driven over the mound of Colony #0056) within a few months prior to the baseline survey in December 2022; a controlled burn of the forest west of the road in the second half of 2024 may have triggered the back-migration in 2024.

**Colony #0095** relocated between January 2023 and January 2024 from a location at a small unpaved forest road into open habitat created by logging in spring 2023 to the south-west of that small forest road, a nest-relocation distance of 32.4 meter. Colony #0095 was first found at its pre-migration site in April 2006; the mound size was recorded as “huge colony” in April 2006, but no estimates of mound dimensions were recorded in 2006. In January 2024, after migration, there were a few active exits on the pre-migration mound (i.e., the pre-migration mound was not completely abandoned), and there were active foraging exits at regular intervals between the pre-migration and post-migration mounds; this indicates that the pre-migration and post-migration mounds were linked by underground tunnels inhabited by the same *A. texana* colony. Colony #0095 remained at its post-migration nest site until at least January 2025. The mound size of Colony #0095 was intermediate (12 meter X 8 meter) in January 2023 at its pre-migration site, increased to 15 meter X 11 meter at its post-migration site by January 2024, and remained approximately the same size (14 meter X 13 meter) in January 2025. The migration was likely triggered by the opening in 2023 of the logged area just west of the pre-migration location of Colony #0095, allowing the colony could migrate out of closed-canopy forest into more open habitat.

**Colony #0097** relocated between January 2024 and January 2025 from the north side of an unpaved forest road to a nearby opening in semi-open forest to the south of that forest road, a nest-relocation distance of 87.3 meter. Colony #0097 was first found at its pre-migration location in April 2006 and was re-surveyed from June 2019 to January 2025. The migration in 2024 was triggered by clear-cut logging of the forest north of that forest road sometime in 2024 (including logging of pine trees surrounding a large forest clearing created by the ant workers for their mound at the side of the forest road). In January 2025, the pre-migration mound site was still heavily disturbed by the logging in 2024 and by the subsequent machine-planting of pine seedlings, but there were active foraging exits next to the pre-migration mound (i.e., the pre-migration site was not completely abandoned), and there were many active foraging exits at regular intervals between the pre-migration mound from January 2024 and post-migration mound from January 2025; this indicates that the pre-migration and post-migration mounds were linked by underground tunnels inhabited by the same *A. texana* colony. Colony #0097 then migrated back between January 2025 and April 2025 to its exact pre-migration location from January 2024, a return distance of 86.7 meter. Colony #0097 is one of the largest *A. texana* colonies included in this study (as estimated by above-ground mound dimensions); its mound size was recorded as "huge mound" in April 2006 (but no estimates of mound dimensions were recorded at that time); 25 meter X 19 meter in June 2019; 37 meter X 20 meter in November 2020; 31 meter X 27 meter in March 2021; 31 meter X 27 meter in January 2023; 28 meter X 24 meter in January 2024 at the post-migration site after the first migration; 31 meter X 27 meter in January 2025 at the post-migration site; and in April 2025, simultaneously 16 meter X 13 meter at its initial pre-migration site and 21 meter X 19 meter at its intermediate migration site (i.e., the colony appeared to occupy two mounds in April 2025 while in the process of the return migration and shifting of work forces between the two nest mounds). The initial leg in 2025 of the back-and-forth migration of Colony #0097 was undoubtedly triggered by the clear-cut logging of the forest north of the forest road in late 2024 (and likely heavy logging machinery driven over the pre-migration mound), but by April 2025 Colony #0097 had shifted part or most of its work force back to its pre-migration location from 2006-2024, after the clear-cut forest north of the forest road had been replanted with new pine seedlings. As described above for other colonies, migrations of colonies into recently re-planted pine plantations with new pine seedlings are not uncommon in *A. texana*.

**Colony #0122** shifted between January 2024 and January 2025 a distance of 21.6 meter along the forest edge south-east of a high-voltage powerline. Colony #0122 was first found at its pre-migration site in November 2020, and Colony #0122 was at that time one of the largest mounds known (29 meter X 21 meter diameter). In January 2024, the mound diameter at the pre-migration site was estimated to be still large (30 meter X 26 meter diameter), but the number of active exits on the mound with fresh excavate seemed reduced; this prompted a search for possible new nest sites nearby to which Colony #0122 may be in the process of migrating. The search revealed a large cluster of fresh excavation crater at the future post-migration site; this large cluster did not appear to be a fully developed mound in January 2024, suggesting that the migration to this satellite mound (i.e., the large cluster of fresh excavation crater) may have started towards the end of 2023, but was still in progress in January 2024. By January 2025, Colony #0122 had relocated to this post-migration mound 21.6 meter distant, and the pre-migration mound seemed more abandoned than occupied: there were a few active exits on and nearby the pre-migration mound (i.e., the pre-migration mound was not completely abandoned), and there were active foraging exits at regular intervals between the pre-migration and post-migration mounds; this indicates that the pre-migration and post-migration mounds were linked by underground tunnels inhabited by the same *A. texana* colony. After migration, the mound size of Colony #0122 was reduced to 13 meter X 12 meter, compared to the pre-migration mound sizes of over approximately 30 meter X 20 meter diameter. The reason for the short shift of 21.6 meter between January 2024 and January 2025 is unclear, as the surrounding forest and the forest edge were not disturbed during this time window, except the roadside between road and forest edge near Colony #0122 was mowed once or twice annually by tractor mowers. The pre-migration mound site had become more and more overgrown with yaupon between 2020-2024, and the colony may have shifted to the new site because the post-migration site was less obstructed by yaupon and trees (half of the post-migration mound was in the open mowed roadside, and half in the shade underneath trees of the forest edge).

**Colony #0149** shifted between January 2023 and January 2024 a distance of 26.4 meter southward along the forest edge west of a paved forest road. Colony #0122 was first found in November 2020, and at that time the pre-migration and the post-migration nest sites were already observed and recorded in the field notes (i.e., the mound was apparently in the process of migration, or polydomous, in January 2023), and the pre-migration mound was judged in November 2020 to be the main mound because it showed overall more fresh excavate on a somewhat smaller mound (19 meter X 14 meter diameter) than the post-migration mound (22 meter X 17 meter dimensions). In January 2023, only the pre-migration mound (24 meter X 23 meter dimensions) was observed and recorded in the field notes (there is no record in our field notes of a live mound at the post-migration site). By January 2024, the colony had shifted from the pre-migration location (15 meter X 7 meter mound dimensions) from January 2023 to the post-migration location (19 meter X 17 meter mound dimensions) from January 2024, there were many excavation crater on both the pre-migration mound and the post-migration mound (i.e., the pre-migration mound was not completely abandoned), but the post-migration mound 26.4 meter distant was judged in January 2024 to be the main mound because it was larger (19 meter X 17 meter mound dimensions) at that time than the pre-migration mound (15 meter X 7 meter). At each survey, there were active foraging exits at regular intervals between the pre-migration and post-migration mounds; this indicates that the pre-migration and post-migration mounds were linked by underground tunnels inhabited by the same *A. texana* colony. Colony #0149 then shifted between January 2024 and January 2025 back northward to its former location (21 meter X 18 meter mound dimensions after back-migration), a back-migration distance of 20.6 meter. The reasons for these two short back-and-forth shifts within 2 years are unclear, as the forest and the roadsides near Colony #0149 were not disturbed during this time window.

**Colony #0180** shifted between January 2023 and January 2024 a distance of 21.7 meter north-eastward from the forest edge at the west side of a gasline cut through pine forest to the forest edge at the east side of that same gasline. Colony #0180 was first found in November 2020, measuring 21 meter X 15 meter mound dimensions. In January 2023, the existence of a cluster of large excavation crater was recorded in the field notes at the future post-migration site (in addition to the main mound of 12 meter X 11 meter diameter at the pre-migration site), so the migration may have already started in January 2023. In January 2024, there were a few active exits on the pre-migration mound (i.e., the pre-migration mound was not completely abandoned), and there were active foraging exits at regular intervals between the pre-migration mound and the large post-migration mound (15 meter X 14 meter); this indicates that the pre-migration and post-migration mounds were linked by underground tunnels inhabited by the same *A. texana* colony. Colony #0180 remained at its post-migration site until at least January 2025, and did not change in mound dimensions (also 15 meter X 14 meter in January 2025). In January 2023, there was a controlled burn of the surrounding forest east and west of the gasline, which may have triggered or accelerated the migration (however, 8 other *A. texana* colonies surveyed within the controlled-burn area did not migrate in response to that controlled burn). The reason for this short shift of 21.7 meter between January 2023 and January 2024 across the gasline therefore is unclear, as the forest and the forest edges near Colony #0180 did not appear to have been disturbed during this time window, except the gasline was mowed annually once by large tractor mowers.

**Migration of Control-Sample Colony between January 2023 – January 2025 during 2-year survey:**

**Colony #0150** shifted between January 2023 and January 2024 a distance of 26.9 meter westward along the forest edge at the north boundary of a young pine plantation. Between the pine plantation and the forest edge was a mowed strip ≈9 meter wide, and Colony #0150 migrated westward within that mowed strip. Colony #0150 was first found in January 2023, and was large (12 meter X 11 meter) in January 2023 at the pre-migration site and somewhat larger (15 meter X 14 meter) in January 2024 at the post-migration site. In January 2024, there were a few active exits on the pre-migration mound (i.e., the pre-migration mound was not completely abandoned), and there were active foraging exits at regular intervals between the pre-migration and post-migration mounds; this indicates that the pre-migration and post-migration mounds were linked by underground tunnels inhabited by the same *A. texana* colony. In January 2025, the main mound of Colony #0150 (16 meter X 15 meter) remained at the post-migration site, but a smaller mound with fresh excavate, recorded as “satellite mound” (13 meter X 13 meter) in the field notes, re-appeared at the pre-migration site. The reason for the short shift of 26.9 meter in 2023 is unclear, as the forest and the forest edges near Colony #0150 did not appear to have been disturbed during this time window, except Colony #0150 was driven-over by tractor mowers cutting the vegetation in the mowed strip at the border of the young pine plantation.

**Summary of the observed 12 colony migrations:**

In eight of the above 12 colony migrations, colonies relocated more than 30 meters; seven of these eight colony migrations of more than 30 meter were linked to physical disturbances, such as logging of trees close to a mound, heavy machinery or trucks driven over a mound, or controlled burns of the forest surrounding a mound (details above). Even though we selected 171 nests in habitat were we thought disturbance by humans was unlikely, some of these nests were impacted by humans during our 2-year survey, most frequently impacted by logging (in locations where we misjudged the age of a pine plantation, and where we wrongly judged logging would not occur during the two years of our survey), and impacted less frequently by controlled burns of forest. It is possible that, in the absence of these unanticipated disturbances such as logging or controlled burns, most or all of the above human-impacted *A. texana* colonies would not have migrated to a new nest site.

A major obstacle of conducting a future, long-term demography study of *A. texana* therefore will be occasional human disturbance that impacts *A. texana* colonies even in protected habitat, such as National Forest.

### NOTES ON *A. TEXANA* COLONIES SCORED DEAD OR MORIBUND IN THE 2-YEAR SURVEY

We summarize here our notes from Table S3 on the history of the 7 *A. texana* colonies that were scored dead and the 2 colonies that were scored moribund in 2025 at the end of our 2-year survey:

**Main Sample, History of Moribund or Dead Colonies in the 2-Year Survey:**

**Colony #0117** (found dead in January 2025, and confirmed dead in April 2025): Colony #0117 was first found in November 2020 in a small clearing in a closed-canopy, managed, pine plantation; the mound of Colony #0117 at that time was estimated to be very large (18 meter X 16 meter). Colony #0117 was re-surveyed in March 2023 (18 meter X 12 meter) and in January 2023 (11 meter X 9 meter); the mound sizes over this time period indicated a decreasing trend from November 2020 to January 2023. In January 2024, no fresh excavation crater were found on the mound of Colony #0117, but in an extensive search, several well-thatched foraging exits were found within a 25-meter radius around the mound. No other foraging exits were found at somewhat greater distances, and no other *A. texana* mound was found when searching a 30-meter circle and an 80-meter circle around the mound (i.e., Colony #0117 did not appear to have migrated). Colony #0117 was therefore scored as alive, but moribund in January 2024. In January 2025, there was no sign of live *A. texana* at the former mound of Colony #0117, and another careful search of the former mound site and its immediate surround, as well as searching a 30-meter circle and an 80-meter circle around the mound, did not reveal any sign of live *A. texana*. Colony #0117 was therefore scored as dead in January 2025. To confirm the dead status of Colony #0117, the former mound site and nearby area was again searched carefully in April 2025; no sign of live *A. texana* was found at the former mound site of Colony #0117 and in the surround, but a small *A. texana* mound (5 meter X 5 meter) was found at 61.0 meter distance at the border of the pine forest at the side of a small forest road. It is possible that Colony #0117 had migrated to that location in an earlier year, but to be conservative, Colony #0117 was scored as dead because no continuous series of foraging exits had been found in earlier searches between the mound site of Colony #0117 and the site of the small mound found later at 61 meter distance. Because Colony #0117 was located in a pine plantation that was not controlled-burned or logged during the 2-year survey, the cause of the death of Colony #0167 is unknown.

**Colony #0136** (found alive but recently poisoned by foresters in January 2024, dead in January 2025, and confirmed dead in April 2025): Colony #0136 was first found in November 2020 at the forest edge to the north of a paved forest road passing through a pine plantation; the mound of Colony #0136 was estimated to be large (14 meter X 9 meter) at that time. The pine forest was judged to be relatively old, but unlikely to be logged within the next few years, so Colony #0136 was included in the Main Sample. At the start of the survey in January 2023, Colony #136 was re-surveyed at the same exact location, and the mound appeared healthy, showed much fresh excavate, and measured 13 meter X 12 meter. A year later in January 2024, Colony #136 was found alive (there were workers excavating at mound exits and foraging exits closeby), but the colony had very recently been poisoned by foresters (a white residue had been applied to many exits, and there was no worker activity at these poisoned exits). The surrounding forest had recently been logged, and the logged forest had not yet been replanted with new pine seedlings by January 2024. Because workers were active at unpoisoned exits on the mound and at unpoisoned foraging exits nearby, and because the colony had obviously been alive 2-3 weeks earlier before the poison was applied, Colony #136 was scored alive for this first resurvey in January 2024. In January 2025, the mound and the surrounding area showed no signs of live *A. texana*, there was much old washed-out excavate at the former mound location, and there were no open exits on this former mound. To confirm the absence of live *A. texana*, a 30-meter circle and an 80-meter circle around the nest was searched carefully; walking these circles was easy because the logged surround had been replanted sometime in 2024 and the new pine seedlings were small in January 2025. Because colony migration was ruled out by the absence of any sign of live *A. texana* within 100 meter of the former mound site, Colony #136 was therefore scored as dead in January 2025. To confirm the absence of live *A. texana*, the search of a 30-meter circle and an 80-meter circle around the nest was repeated in April 2025, and again no signs of live *A. texana* were found at the former mound site and in the surround of this mound. Although Colony #0136 did not die naturally but was poisoned by foresters, Colony #0136 was scored as dead in the final Main Sample, inflating our estimate of natural mortality of *A. texana* in "stable" habitat where we had expected infrequent impact by humans. The cause of death of Colony #136 was undoubtedly poisoning by foresters.

**Colony #0167** (found moribund in January 2024, dead in January 2025, and confirmed dead in April 2025): Colony #0167 was first found in January 2023 in semi-open pine forest at the side of the Backbone Trail in the Kisatchie National Forest. The mound of Colony #0167 at that time was estimated to be relatively small (7 meter X 5 meter), with fresh excavation crater (some of them well-thatched), but with many leaves on the mound. The presence of many leaves on the mound could indicate that Colony #0167 was already declining in 2023, because leaves fallen onto an established healthy mound become buried typically by newly-excavated soil discarded by workers on the mound. In January 2024, there were no excavation crater with fresh excavate on the mound site, but there were three foraging exits with workers within 25 meter distance from the mound site. There were no other signs of *A. texana* when searching a 30-meter circle and an 80-meter circle around the former nest site, indicating that Colony #0167 had not migrated to a location nearby, but that the colony was moribund. In January 2025, the former nest site was covered completely with a thick layer of leaves, and there were no signs of live *A. texana* at and near the former nest site. We repeated the search of a 30-meter circle and an 80-meter circle around the former nest site, and the absence of any signs of live *A. texana* indicated that Colony #0167 had died. Because reaching Colony #0167 requires a time-consuming hike, we did not confirm the death of Colony #0167 by another re-survey in April 2025 (as we did in April 2025 for the other moribund and dead colonies found in our 2-year survey). However, we were able to re-survey the former nest site of Colony #0167 in March 2026, and in a quick search we found no signs of live *A. texana* at or near the former nest site of Colony #0167. Given this history, we scored Colony #0167 as moribund in January 2024, and as dead in January 2025. Because Colony #0167 was located in a remote area of National Forest that was not controlled-burned during the 2-year survey (the forest was controlled-burned in winter 2026 after completion of our 2-year survey), the death of Colony #0167 was presumably due to natural causes, not due to human harm. The actual cause of the death of Colony #0167 is unknown.

**Colony #0058** (found moribund in January 2025, and confirmed moribund in April 2025): Colony #00058 was first found in November 2020 at the forest edge to the east of a paved road passing through managed pine forest (forest that is regularly controlled-burned, but not logged); the mound of Colony #00058 at that time was estimated to be very large (17 meter X 15 meter). Colony #00058 measured 12 meter X 11 meter in December 2022 at the start of our 2-year survey (most nests at the start of the 2-year survey were surveyed in January 2023, but some nests, like Colony #00058, were surveyed a month earlier in December 2022); the mound size was somewhat difficult to estimate at that time because many pine needles had accumulated on the mound. In January 2024, there were fresh excavation crater on the mound (because of oversight, the size of the mound was accidentally not recorded at that time), and there were several foraging exits with workers within 40 meter radius of the nest site; Colony #00058 was therefore scored as alive, but possibly declining. A search of a 20-meter circle and a 80-meter circle around the nest site did not suggest the presence of another large mound nearby (the closest large live *A. texana* colony, Colony #00059, was 153 meter distant; because of this significant distance, it is unlikely that Colony #00058 and Colony #00059 were linked via underground tunnels, but it is possible). In January 2025, many large branches and much pine bark had fallen onto the mound because of a major storm causing a treefall closeby (so it was difficult to estimate the size of the mound), some fresh excavate was hidden under the fallen branches, and there were active foraging exits with workers within 15 meter distance from the mound. No other *A. texana* mound was found when searching a 30-meter circle and an 80-meter circle around the mound. Because Colony #00058 seemed already declining in January 2024, did not recover by January 2025, and seemed again noticeably smaller compared to the mound sizes of Colony #00058 recorded in the surveys in 2020 and 2022, we scored Colony #00058 conservatively as moribund in January 2025. To confirm this status of moribund, we revisited the mound of Colony #00058 in April 2025; we found at that time four fresh excavation crater with workers hidden under the fallen branches, and we found one foraging exit with workers within 15 meter distance from the mound, confirming that a score of moribund was more appropriate for Colony #00058 in early 2025 than a score of alive and healthy (i.e., not declining). The cause of the morbidity of Colony #00058 is unknown.

**Colony #0164** (found moribund in January 2025, and confirmed moribund in April 2025): Colony #0164 was first found in November 2020 at the forest edge to the west of an unpaved county road passing through mixed oak-pine forest (likely unmanaged and not controlled-burned forest); the mound of Colony #0164 at that time was estimated to be large (13 meter X 12 meter). Colony #0164 measured 10 meter X 9 meter in December 2022 at the start of our 2-year survey (most nests at the start of the 2-year survey were surveyed in January 2023, but some nests, like Colony #0164, were surveyed a month earlier in December 2022), and the colony appeared vigorous at that time. In January 2024, the mound was covered with many pine needles and was therefore difficult to size, but the presence of workers on the mound, as well as the presence of nearby active foraging exits, indicated that the colony was alive. In January 2025, a tree trunk had fallen onto the mound, and we found no fresh excavate and observed no workers on the mound, but we found foraging exits closeby. No other *A. texana* mound was found when searching a 30-meter circle and an 80-meter circle around the mound, so we scored Colony #0164 conservatively as moribund. To confirm this status of moribund, we revisited Colony #0164 in April 2025; we found again excavate hidden under the tree/branches that had fallen onto the mound, saw no workers on the mound, but we found a foraging exit about 15 meter distant from the mound. These observations confirmed that a score of moribund was more appropriate for Colony #0164 in early 2025 than a score of alive and healthy (i.e., not declining). The cause of the morbidity of Colony #0164 is unknown.

**Control Sample, History of Moribund or Dead Colonies in the 2-Year Survey:**

**Colony #0007** (found dead in January 2025, and confirmed dead in April 2025): Colony #0007 was first found in November 2020 at the west-border fence of a cemetery; the mound of Colony #0007 at that time was estimated to be large (12 meter X 7 meter). Half of the mound was inside the cemetery (i.e. the mound center was underneath the fence), and that half had recently been mowed over by cemetery caretakers mowing the cemetery lawns. Colony #0007 measured 9 meter X 8 meter in December 2022 at the start of our 2-year survey (most nests at the start of the 2-year survey were surveyed in January 2023, but some nests, like Colony #0007, were surveyed a month earlier in December 2022); the mound had shifted by that time ≈10 meter westward to the forest edge just outside the cemetery. Because Colony #0007 had many active foraging exits among the graves throughout the western part of the cemetery, we included this nest in the Control Sample (possible disturbance by humans, because nests in cemeteries are sometimes poisoned by cemetery caretakers). In January 2024, the mound measured 12 meter X 9 meter, appeared healthy and vigorous as before, and was located at the same site at the forest edge just outside the cemetery. As in earlier years, the nest had many active foraging exits among the graves throughout the western part of the cemetery. In January 2025, there was no sign of live *A. texana* in the western part of the cemetery (no excavate on the former mound, no active foraging exits nearby), and a 10-meter-wide strip of forest just outside the cemetery fence had been cleared within the last year, including forest right at the nest site of Colony #0007. No other *A. texana* mound was found in January 2025 when searching a 30-meter circle and an 80-meter circle through forest and cemetery around the mound. Because of the absence of any sign of live *A. texana* within 100 meter of the location of the former mound of Colony #0007, we scored Colony #0007 as dead in January 2025. In April 2025, we resurveyed the cemetery and found again no sign of live *A. texana* within 100 meter of the former mound of Colony #0007. A caretaker mowing the lawns in the cemetery that day stated that the leafcutter mound at the fence was poisoned in 2024 with “white poison” when the forest strip just to the outside of the cemetery was cleared.

**Colony #0078** (found dead in January 2025, and confirmed dead in April 2025): Colony #0078 was first found in July 2019 right at the roadside to the west of a paved country road; the mound of Colony #0078 extended likely also under the road, and the overall mound size was therefore difficult to estimate (no mound size was recorded at that time). Google Street View shows this mound having at least 10-meter diameter in March 2008, at the exact location where we found this mound in 2019, at the west side of the road, and extending under the road to the east roadside. Google Street View shows that the forest to the west was mature in 2008. By 2019, the forest to the west had been logged and replanted within a few years before 2019. In January 2020, the mound of Colony #0078 measured 7 meter X 6 meter (plus a nest extension under road that was not visible from the paved surface). At the start of our survey in January 2023, the mound measured 8 meter X 4 meter (plus extension under road). Because *A. texana* nests built by the ants at roadsides and under roads can cause the pavement to crack or sag as the ants excavate large garden cavities under the pavement (sagging and cracking of pavement is very typical for *A. texana* nests tunneling by *A. texana* ants under paved roads), some road maintenance crews prevent road erosion by poisoning leafcutter nests located right at roadsides; we therefore included Colony #0078 in the Control Sample (possible disturbance by humans, specifically possible poisoning by road maintenance crew to prevent road erosion). Between January 2023 and January 2024, the highway department repaved a large patch at the roadside of the county road right where Colony #0078 was located (this repaved patch is visible in a Google Street View image from November 2024), presumably because of road erosion caused by the colony's excavation of soil from underneath the road. It is possible that during the re-paving the highway department poisoned Colony #0078, because in January 2024, we found the size of the mound greatly reduced compared to January 2023. Specifically, we scored Colony #0078 as moribund for January 2024 because we found only 4 exits with workers on the mound (no fresh excavate on the mound) and because we did not find other signs of live *A. texana* when searching a 30-meter (radius) and an 80-meter circle around the mound (also, there were no fresh excavation crater at the other roadside, which is typical for large nests built underneath a paved road), indicating that that Colony #0078 had not migrated to somewhere nearby. In January 2025, there was no sign of live *A. texana* at the former mound location of Colony #0078, and again we did not find other signs of live *A. texana* when searching a 30-meter (radius) and an 80-meter circle around the mound, so we scored Colony #0078 as dead. To confirm the status of dead, we re-surveyed the former mound site of Colony #0078 in April 2025, and again failed to find live *A. texana* at the former mound site and nearby. Although it is possible that #0078 was poisoned because it had constructed its nest partly under a paved county road, the cause of death of #0078 is unknown.

**Colony #0131** (found dead in January 2025, and confirmed dead in April 2025): Colony #0131 was first found in November 2020 in a protected, unused corner of a cemetery, but with graves within ≈30 meter distance; the size of the mound of Colony #0131 was difficult to estimate at that time because many oak leaves had recently fallen onto the mound. Colony #0131 had many large fresh foraging exits among the graves throughout the cemetery. At the start of our survey in January 2023, the mound was exceptionally large, measured 22 meter X 16 meter, and showed very much fresh excavate. Because of the nest's many active foraging exits among the graves throughout the cemetery, we included this nest in the Control Sample (possible disturbance by humans, because nests in cemeteries are sometimes poisoned by cemetery caretakers). Shortly before the survey day in January 2024, cemetery caretakers had raked branches and leaves onto the mound of Colony #0131 to start a major fire there, and by the day of the survey, the workers of Colony #0131 had managed to dig out at only a few exits. A major fire burning for hours on a mound typically does not kill an *A. texana* colony, but because we found only a few active excavation crater at the mound site, we scored Colony #0131 conservatively as moribund for January 2024. In January 2025, we found only old excavate at the former mound site of Colony #0131, and no active foraging exits throughout the cemetery and when searching a 30-meter (radius) and an 80-meter circle around the mound, so we scored Colony #0078 as dead. To confirm the status of dead, we re-surveyed the former mound site of Colony #0131 in April 2025, and again failed to find signs of live *A. texana* at the former mound site and nearby. It is possible that the cemetery caretakers first tried to kill the *A. texana* colony by building a major fire on the mound in January 2024, then may have poisoned the colony sometime later in 2024 because the fire did not killed the colony completely. The cause of death of Colony #0131 appears to have been human-induced harm (fire, possible poison).

**Colony #0152** (found dead in January 2025, and confirmed dead in April 2025): Colony #0152 was first found in November 2020 right at the roadside of, and partly underneath, a narrow paved county road passing through an older pine plantation; the mound of Colony #0152 at that time was estimated to be relatively small (8 meter X 7 meter, plus a nest extension under the paved road that was not visible from the surface). In January 2023 at the start of our survey, the mound measured 15 meter X 10 meter between roadside and forest edge, but the mound was difficult to size accurately because there were many leaves on the mound. Because Colony #0152 was located right next to a paved road and the nest was partly underneath that road, we included this nest in the Control Sample (possible disturbance by humans, because nests under paved roads are sometimes poisoned by road maintenance crews). In January 2024, Colony #0152 appeared healthy and vigorous (much excavate on the mound); the mound measured 14 meter X 8 meter, but the mound was again difficult to size because of many leaves on the mound. In January 2025, we found no sign of live *A. texana* on the mound and also when searching a 30-meter (radius) circle around the mound, so we scored Colony #0152 as dead. To confirm the status of dead, we re-surveyed the former mound site of Colony #0152 in April 2025, and again failed to find signs of live *A. texana* at the former mound site and nearby. Although it is possible that Colony #0152 was poisoned because it had constructed its nest partly under a paved road, the cause of death of Colony #0152 is unknown.

**Summary of the 7 dead and 2 moribund colonies scored in January 2025 at end of 2-year survey**

| **COLONY ID** | **LIKELY CAUSE OF DEATH OR OF MORBIDITY** |
|---|---|
| **Main Sample** | |
| Colony #0117 | cause of death unknown |
| Colony #0136 | poisoning by foresters after logging of the surrounding pine plantation |
| Colony #0167 | cause of death unknown |
| Colony #0058 | cause of morbidity unknown |
| Colony #0164 | cause of morbidity unknown |
| **Control Sample** | |
| Colony #0007 | killed by cemetery caretakers |
| Colony #0078 | possibly killed by road maintenance crew, but cause of death unknown |
| Colony #0131 | likely killed by cemetery caretakers |
| Colony #0152 | cause of death unknown |
